# Parvovirus infection affects host cell nucleolar organization and ribosome biogenesis

**DOI:** 10.1101/2022.06.07.495090

**Authors:** Moona Huttunen, Satu Hakanen, Vesa Aho, Simon Leclerc, Aynur Soenmez, Sergey Kapishnikov, Alessandro Zannotti, Visa Ruokolainen, Kari Salokas, Markku Varjosalo, Leena Latonen, Colin R Parrish, Kenneth Fahy, Salla Mattola, Denis L.J. Lafontaine, Maija Vihinen-Ranta

## Abstract

The nucleolus is a biomolecular condensate essential for ribosome biogenesis and cellular stress response, and it is a key target for many DNA viruses. However, little is known about how autonomous parvovirus infection impacts nucleolar structure and function. Here, we used advanced imaging techniques, including ten-fold robust expansion microscopy (TREx), cryo soft X-ray tomography (Cryo-SXT), and interactomics and biochemical approaches, to study nucleolar remodeling during canine parvovirus infection. Infection led to redistribution of nucleolar upstream binding transcription factor 1 (inner core), fibrillarin (middle layer), and Ki-67 (outer rim). In contrast, peripheral nucleolar proteins (nucleolin and nucleophosmin) and precursor ribosomal RNAs (pre-rRNAs) remain in circularized structures. TREx and Cryo-SXT microscopy revealed profound nucleolar structural changes, including thickened perinucleolar chromatin and enlarged nucleolar low-protein density channels. BioID identified interactions between viral NS2 and nucleolar proteins in ribosome biogenesis. Northern blotting demonstrated a slowdown in ribosome biogenesis during infection. Collectively, these findings provide novel insights into how parvoviruses remodel nucleolar structure and function.

## Introduction

Interaction with a multifunctional subnuclear compartment, the nucleolus, is crucial for the replication and pathogenesis of several DNA viruses, including herpesviruses (Boyne and Whitehouse, 2009; Pearson et al., 2011) and adenoviruses (Lee et al., 2003; Hindley et al., 2007; Lawrence et al., 2006). Moreover, the capsids of adeno-associated virus (AAV), a helper-dependent parvovirus, are assembled in the nucleolus (Wistuba et al., 1997; Johnson and Samulski, 2009). The role of nucleoli in the autonomous parvovirus life cycle has remained unexplored.

The nucleolus is the largest nuclear subcompartment, and it is involved in the initial steps of ribosome biogenesis, such as transcription of ribosomal RNA genes (rRNA), processing and modification of precursor rRNAs (pre-rRNAs), and ribosome subunit assembly (Dubois and Boisvert, 2016; Raska et al., 2006). The events of ribosome production are most likely coordinated, at least in part, by liquid-liquid phase separation (Lafontaine, 2019; Yao et al., 2019). The nucleolus is surrounded by heterochromatin and euchromatin, forming a stable nucleolus-nucleoplasm interface around the nucleoli (Qi and Zhang, 2021; Padeken and Heun, 2014; Caragine et al., 2019; Peng et al., 2023). The mammalian nucleolus comprises three main morphological components: the fibrillar center (FC), the dense fibrillar component (DFC), and the granular component (GC) (Pederson, 2011; Hernandez-Verdun, 2011; Raska et al., 2006; Shan et al., 2023; Lafontaine, 2023a; b). Additional subphases have recently been identified by high-resolution fluorescence microscopy, including the periphery of the dense fibrillar center (PDFC) and the nucleolar rim (NR) (Shan et al., 2023; Stenström et al., 2020). FCs and DFCs are involved in rDNA transcription and initial maturation of pre-rRNAs, whereas further processing of pre-rRNAs and assembly of ribosome subunits occur in GC (Pederson, 2011; Hernandez-Verdun, 2011; Raska et al., 2006). The nucleolus is a dynamic structure (Feric et al., 2016; Falahati and Wieschaus, 2017; Lafontaine et al., 2021; Strom and Brangwynne, 2019) with highly mobile proteins constantly shuttling between the nucleolus and the nucleoplasm (Phair and Misteli, 2000; Andersen et al., 2005). The nucleolus has many additional functions besides ribosomal biogenesis, including the regulation of mitosis and the cell cycle and acting as a biosensor of stresses of different origins, such as oxidative stress, heat shock, nutrient deprivation, and viral infection (Boisvert et al., 2007; Iarovaia et al., 2019; Boulon et al., 2010; Scherl et al., 2002; Salvetti and Greco, 2014).

The proper localization of several nucleolar proteins is profoundly affected by viral infection. In healthy cells, upstream-binding factor 1 (UBF1), fibrillarin (FBL), nucleolin (NCL), nucleophosmin 1 (NPM1), and proliferation marker protein Ki-67 are enriched in the nucleolar compartments FC, DFC, PDFC, GC, and NR, respectively (Lafontaine et al., 2021; Stenström et al., 2020). In infected cells, some of these proteins are displaced and even relocated outside of the organelle (Callé et al., 2008; Lymberopoulos and Pearson, 2007; Greco et al., 2012). One remarkable example is the abundant multifunctional phosphoprotein NCL (Biggiogera et al., 1990; Ginisty et al., 1999; Srivastava and Pollard, 1999), which relocalizes from the nucleolus to the viral replication compartments (VRC) upon infection with herpes simplex virus type 1 (HSV-1) (Callé et al., 2008; Lymberopoulos and Pearson, 2007; Greco et al., 2012) and other stresses (Jia et al., 2017; Daniely and Borowiec, 2000). NCL is a multifunctional factor implicated in virtually all stages of ribosome biogenesis (Ginisty et al., 1999; Srivastava and Pollard, 1999), requiring its dynamic shuttling between the nucleolus and the nucleoplasm (Chen and Huang, 2001). Other nucleolar proteins whose distribution is sensitive to stress include FBL, UBF1, and NPM1 (Galibert et al., 2001; Yang et al., 2016; Yao et al., 2010). UBF1 is a transcription factor that binds specifically to the rDNA promoters and maintains the active state of rDNA (Roussel et al., 1993; Tuan et al., 1999; Sanij and Hannan, 2009; Sanij et al., 2008). FBL is a snoRNA-guided methyltransferase responsible for rRNA 2’-O methylation and a pre-rRNA-processing factor essential for the initial steps of ribosomal biogenesis (Lafontaine and Tollervey, 2000; Omer et al., 2002; Tollervey et al., 1993). In addition, FBL binds to nascent transcripts, helping their transition from the FC/DFC interphase to the DFC, where they undergo initial maturation (Lafontaine, 2019). NPM1 assists in ribosome assembly, chromatin modeling, and cell homeostasis (Lindström, 2011). NPM1 is a pentameric protein that mediates the assembly of the GC via heterotypic interactions with nucleolar proteins and rRNA and homotypic interactions with intrinsically disordered regions of nucleolar proteins (Mitrea et al., 2016, 2018). Ki-67 is a “chromosome surfactant” protein located in the nucleoli in interphase cells and the mitotic chromosome periphery in mitotic cells (Verheijen et al., 1989; Kill, 1996). During the interphase, Ki-67 has a specific role in the organization of heterochromatin (Sun and Kaufman, 2018; Sobecki et al., 2016; Stenström et al., 2020); during mitosis, it is essential to reassemble the nucleolus (Lafontaine et al., 2021; Stenström et al., 2020).

Autonomous parvoviruses have non-enveloped icosahedral particles of ∼25 nm in diameter, enclosing a linear single-stranded DNA genome of ∼5.0 kb. A typical parvovirus genome contains two open reading frames that encode several structural and non-structural proteins (Cotmore et al., 1983). The canine parvovirus (CPV) capsid is composed of the structural viral proteins VP1 and VP2, with the latter also being N-terminally cleaved into a third structural protein, VP3, in mature virions (Weichert et al., 1998). CPV and other protoparvoviruses encode two non-structural proteins, NS1 and NS2 (Cotmore et al., 1983). The CPV NS1 protein is a multifunctional protein that exhibits site-specific DNA binding, ATPase, helicase, and nickase activities (Niskanen et al., 2010, 2013). NS1localizes into the nuclear VRC distributed throughout the nucleus but is absent from the nucleolar region (Ihalainen et al., 2009). NS2 of another protoparvovirus, the minute virus of mice, has been linked to viral DNA replication, translation of viral mRNA, capsid assembly, and pre-lytic nuclear egress (Cotmore et al., 1997; Naeger et al., 1990; Eichwald et al., 2002). Although the function of CPV NS2 is currently mostly undefined, our recent studies have revealed a potential role in chromatin remodeling and DNA damage response during replication (Mattola et al., 2022b).

Here, we investigated how CPV infection impacts nucleolar morphology and function. We show that infection leads to the redistribution of key nucleolar proteins and rRNA, extensive remodeling of nucleolar structure, and reorganization of nucleolus-associated chromatin. Notably, these alterations are accompanied by marked changes in pre-rRNA processing, highlighting the profound effects of CPV on nucleolar homeostasis.

## Results

### Nucleolar structure is remodeled during infection

To test if viral infection impacts nucleolus homeostasis and ribosome biogenesis, we analyzed the distribution of critical nucleolar proteins and rRNA in CPV-permissive Norden Laboratory Feline Kidney (NLFK) cells. We examined the distribution of representative markers of the major nucleolar compartments. We used UBF1, FBL, NCL, NPM1, and Ki-67 antibodies to label FC, DFC, PDFC, GC, and NR, respectively (**Fig. 1 a**). NS1 detection was used as a proxy to follow the progression of CPV infection.

**Figure 1.**
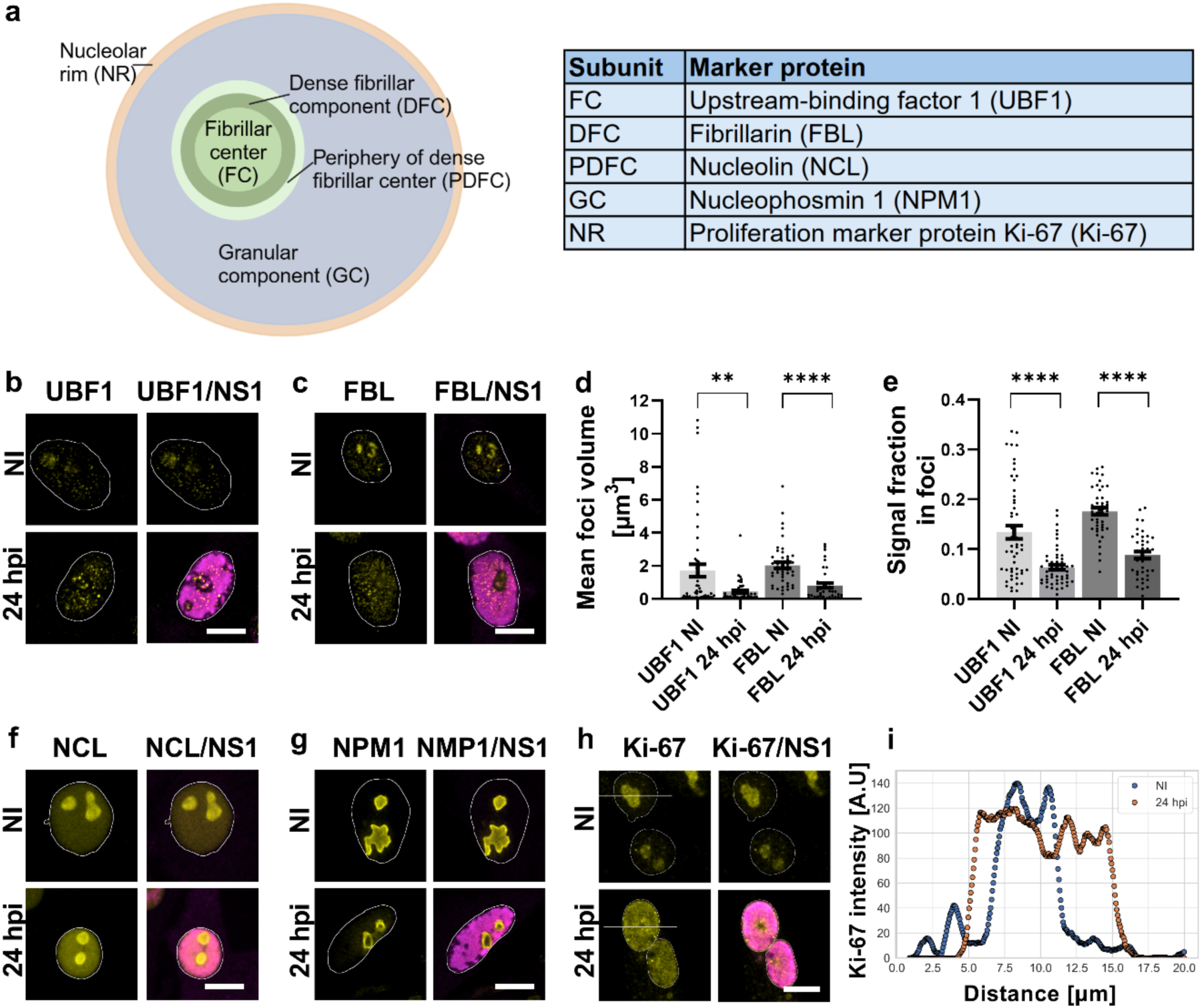
Nucleolar protein distribution changes during CPV infection. (**a)** Schematic of distribution of nucleolar proteins upstream-binding factor 1 (UBF1), fibrillarin (FBL), nucleolin (NCL), nucleophosmin 1 (NPM1), and Ki-67 in nucleolar subcompartments: the fibrillar center (FC), the dense fibrillar component (DFC), the periphery ofthe dense fibrillar center (PDFC), the granular component (GC), and the nucleolar rim (NR). Representative confocal maximum intensity projections showing the distribution of(**b**) UBF1 and (**c**) FBL in noninfected (NI) and infected NLFK cells at 24 hpi. (**d**) Quantitative analyses of the mean volume ofUBF1 and FBL foci, and (**e**) the fraction of nuclear UBF1 and FBL signal detected inthe foci in noninfected and infected cells at24 hpi (n_NI_= 52, 46 and n_inf_= 52, 40, respectively). Distribution of(**f**) NCL, (**g**) NPM1, and (**h**) Ki-67 in noninfected and infected cells at 24 hpi. (**i**) Representative Ki-67 intensity line profiles of noninfected (blue) and infected cells (orange) at 24 hpi. The cells were immunolabeled with antibodies against UBF1, FBL, NCL, NPM1, Ki-67 (yellow), and the viral NS1 protein (a marker of viral replication compartment region, magenta) antibodies. DNA was labeled with DAPI, and a white line illustrates the borders of the nuclei. Scale bar, 10 µm. The error bars show the standard error of the mean. Statistical significance was determined using Welch’s *t*-test. The significance values shown are denoted as **** (p<0.0001) or** (p=0.0021).

At 24 hours post infection (hpi), UBF1 and FBL were relocalized outside of the nucleolar region forming small, punctate foci scattered throughout the nucleoplasmic space while retaining some residual localization in the nucleolus. The emergence of small viral NS1-positive foci, visible at 8 hours post-infection (hpi), initiated the formation of the VRC (**Fig. S1)**. The expansion and coalescence of these foci eventually resulted in an enlarged VRC filling the entire nucleoplasm (**Fig. 1 b and c, Fig. S1 and S2**). Mean volume analysis confirmed that the average size of UBF1 and FBL foci within the nuclear area decreased as the infection progressed up to 24 hpi (**Fig. 1 d**). This was accompanied by a reduction in the fraction of nuclear UBF1 and FBL signal detected in these foci (**Fig. 1 e**), favoring a more diffuse staining pattern. Remarkably, the behavior of PDFC and GC proteins was distinctly different. NCL and NPM1 retained their association with nucleoli during the progression of infection. However, the roundness of NCL and NPM1 regions increased at late infection, indicating a change in phase behavior (**Fig. 1 f and g, Fig. S3 a and S4**). The measurement of NCL-stained regions suggested that the volume of the nucleoli remained unchanged in infection (**Fig. S3 b)**. Finally, Ki-67 representing the NR regions was relocalized from the nucleoli into foci scattered throughout the nucleoplasm in infected cells **(Fig. 1 h and i).** The infection led to a partial Ki-67 redistribution from the nucleoli to the nucleoplasm at 8 hpi. At 12 hpi, Ki-67 was seen in the nucleoli and nucleoplasm, accumulating close to the NE, which likely correlates with the reorganization of the host cell chromatin close to the nuclear envelope as the infection progresses **(Fig. S5, and** see also below **Fig. 3 a**). Later, at 24 hpi, Ki-67 showed a homogenous nuclear distribution with a few distinct small foci **(Fig. S5)**.

Quantitative 3D image analysis demonstrated that the volume of rRNA-containing nucleolar regions decreased (**Fig. 2 a and b**). At the same time, the nucleolar intensity of rRNA remained unchanged at late infection compared to control cells (**Fig. 2 c**). Importantly, similar to the observations with the NCL and NMP1 (**Fig. 1 f and g, Fig. S3 a and S4**), rRNA-containing nucleolar regions became increasingly circular in late infection at 24 hpi (**Fig. 2 d**).

**Figure 2.**
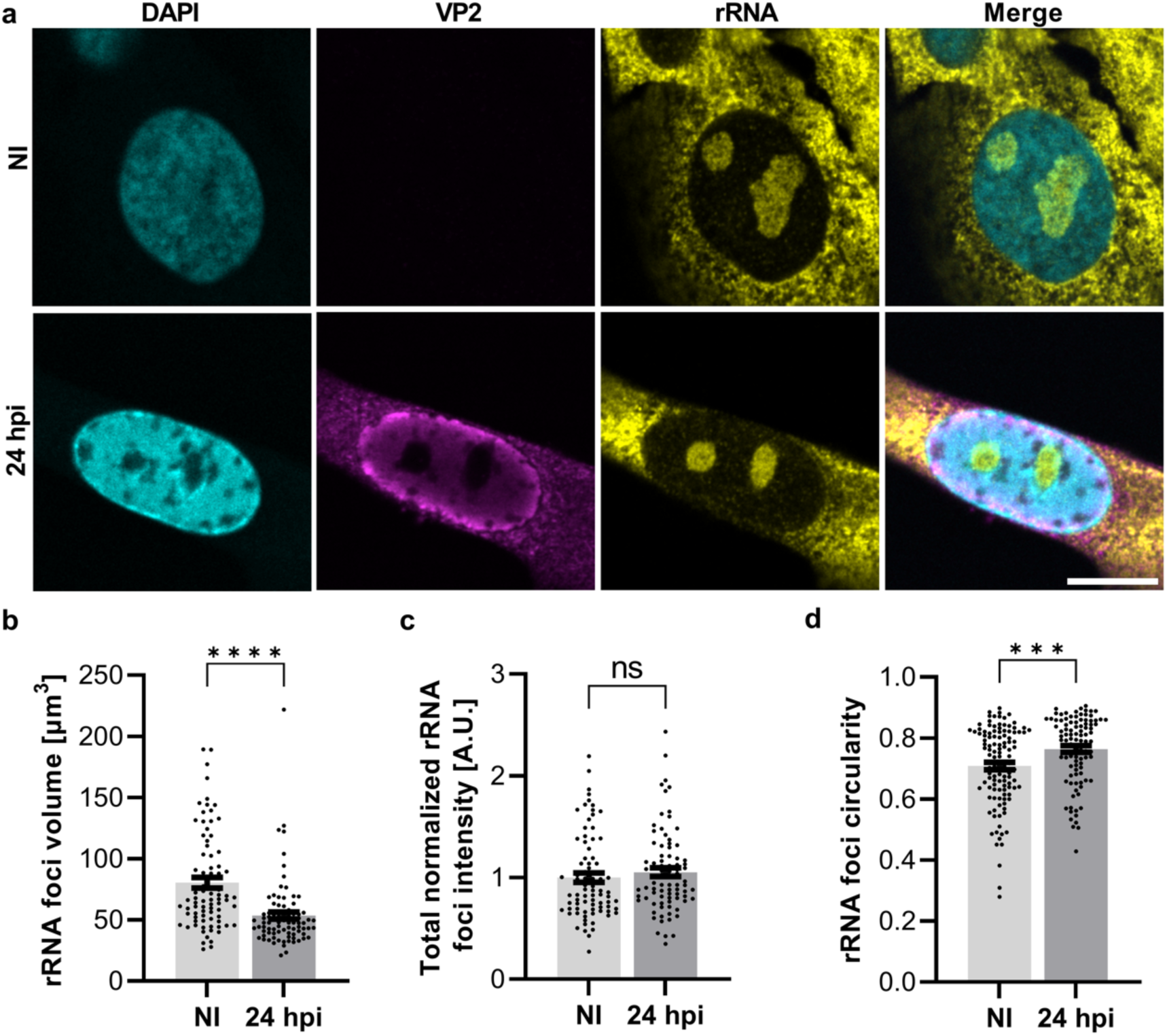
Infection decreases the size and increases the circularity of nucleolar rRNA-positive regions. (**a**) The representative confocal images show the distribution of viral capsids and rRNA in noninfected and infected NLFK cells at 24 hpi. The cells were stained with the viral capsid protein VP2 (magenta) and rRNA (yellow) antibodies. The chromatin was labeled with DAPI (cyan). Scale bar, 10 µm. (b) The mean volume, and (c) the total normalized intensity of segmented rRNA foci in noninfected and infected cells (n = 82 and 84, respectively). (d) The circularity of the rRNA-labeled segmented nucleoli cross-sections in noninfected and infected cells at 24 hpi (n =115 and 100, respectively). The error bars show the standard error of the mean. Statistical significance was determined using Welch’s *t*-test. The significance values shown are denoted as **** (p<0.0001), *** (p=0.0009), or ns (not significant).

In conclusion, upon viral infection, the nucleolus appears to disassemble, starting from its inner core (FC and DFC), which loses integrity, while the more peripheral layers (PDFC and GC) remain somewhat organized but exhibit increased circularity. The NR, which lines the exterior ofthe nucleolus, is also lost.

### Reorganization of nucleolar ultrastructure upon infection

Host chromatin displacement and marginalization near the nuclear envelope (NE) is observed at the late stages of parvovirus infection as the VRC expands (Ihalainen et al., 2009). Little is known about how the infection and infection-induced redistribution of chromatin affect the organization of nucleolus-associated chromatin.

We used ten-fold robust expansion microscopy (TREx) to examine nucleolar chromatin organization and overall protein distribution. TREx enables super-resolution imaging, approaching 25-30 nm resolution, using a conventional confocal microscope by embedding immunostained cell samples in a swellable hydrogel (Damstra et al., 2022). Labeling total protein with a fluorescent NHS ester dye reveals the nanoscale organization and protein intensity within nucleoli, while DAPI staining highlights chromatin distribution (**Fig. 3 a**) (Sheard et al., 2023).

**Figure 3.**
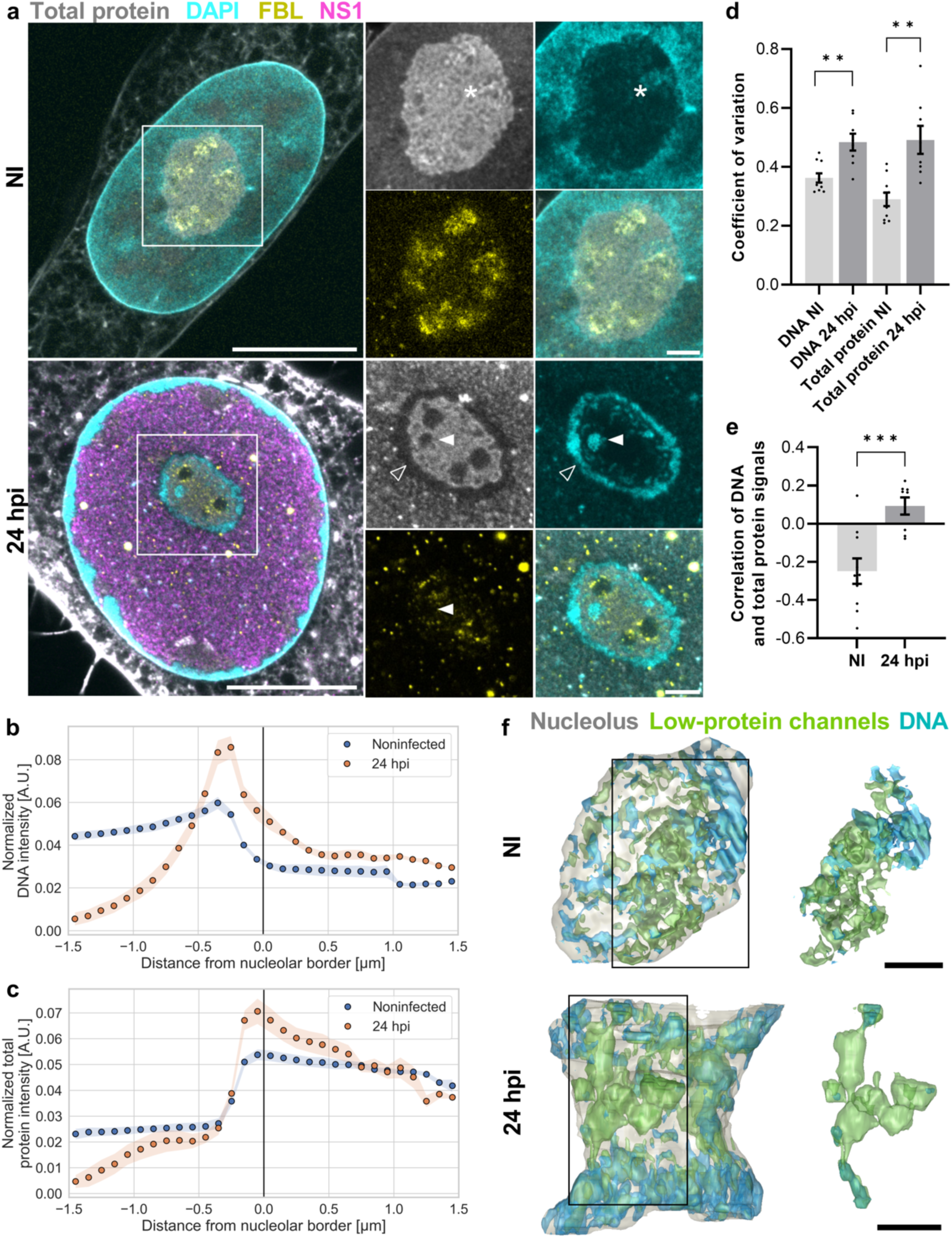
Intranucleolar ultrastructure and chromatin distribution are remodeled in infection. (**a**) Visualization of the nucleolar structure by ten-fold robust expansion microscopy (TREx) in noninfected (NI) and infected NLFK cells at 24 hpi. The representative images show nuclear and nucleolar ultrastructures with the NHS ester total protein stain (gray), DNA staining by DAPI (cyan), and fibrillarin and viral NS1 staining by antibodies (yellow and magenta, respectively). The light areas in total protein-stained cells indicate the presence of highly concentrated protein, and dark areas indicate relatively small amounts of proteins. The final expansion was estimated to be ∼7.5-fold. The localization of nucleolar chromatin in noninfected cells (asterisks), perinucleolar chromatin (empty arrowheads), and nucleolar chromatin (filled arrowheads) in infected cells are shown. The intensities in each image have been independently adjusted for visualization purposes. Scale bars (adjusted to expansion factor), 5 µm and 1 µm. The mean normalized intensity of(**b**) DAPI-labeled DNA and (**c**) total protein as a function of the distance from the nucleolar border in noninfected and infected cells at 24 hpi (n = 10 and 8, respectively). (**d**) The distribution and dispersion of nucleolar DNA and total protein were measured using a coefficient of variation in noninfected and infected cells at 24 hpi. (**e**) DNA and total protein signal correlation in the nucleoli of noninfected and infected cells at 24 hpi (**f**) 3D reconstructions of TREx data show low-protein regions (green) and DAPI DNA stain (blue) in the nucleoli (grey) of noninfected and infected cells. Images on the right depict a single continuous low-protein channel network. Scale bar (adjusted to expansion factor), 1 µm. See also Supplementary Movie 1. The nucleolar segmentation was based on the total protein staining. The error bars show the standard error of the mean. Statistical significance was determined using Welch’s *t*-test. The significance values shown are denoted as ***(p=0.0007) or ** (p=0.0034, p= 0.0033).

The NHS ester labeling revealed protein accumulation in spherical, nucleolus-like regions within the central nuclear areas of both noninfected and infected NLFK cells at 24 hpi. Immunostaining for FBL showed that, in noninfected cells, FBL localized to protein-dense areas – likely corresponding to DFCs – as expected for typical nucleolar architecture (**Fig. 3 a**). Consistent with earlier observations (**Fig.1cande**), TRExmicroscopy at24hpi confirmed partial redistribution of FBL into nucleoplasmic foci upon infection (**Fig. 3 a**), in agreement with disassembly of the FCs and DFCs, previously inferred from the nucleolar release of UBF1 and FBL (**Fig. 2 a**).

Analysis of DNA and total protein distribution revealed that in both noninfected and infected cells, chromatin remained condensed near the nuclear envelope and partially around nucleoli (**Fig. 3 a and b**). However, perinucleolar chromatin intensity was markedly higher in infected cells, coinciding with reduced protein signal in these chromatin-dense zones (**Fig. 3 a and c**). In contrast, regions just inside the nucleolar border showed increased protein intensity during infection (**Fig. 3 c**). The infection also led to a more irregular distribution of DNA within nucleoli, with the formation of large intra-nucleolar DNA foci and increased variability compared to noninfected cells (**Fig. 3 a, d)**. These foci coincided with areas of low protein density and were often surrounded by FBL, yet did not overlap with condensed perinucleolar DNA (**Fig. 3 a,** white arrowhead at 24 hpi). This pattern suggests that the infection drives DNA accumulation in nucleolar interstices, as previously identified by EM studies (Thiry, 1993; Thiry and Lafontaine, 2005; Lafontaine et al., 2021). Finally, infection also disrupted the spatial relationship between DNA and protein: whereas DNA and total protein signals were largely anti-correlated in noninfected cells, they became positively correlated upon infection (**Fig. 3 d, e**).

The 3D reconstructions of TREx data demonstrated that the low-intensity protein regions formed interconnected channel networks, which sometimes spread throughout the nucleoli in noninfected and infected cells. In infected cells, the channels formed larger spherical vacuolar cavities, which were also interconnected. Notably, a substantial part of channels in noninfected and infected cells were DNA-containing interstices, or DNA was located close to the channels (**Fig. 3 f and movie M1**).

To further examine the infection-induced changes in the 3D structure of nucleoli, NLFK cells were grown and infected on electron microscopy (EM) grids and analyzed using cryo soft X-ray tomography (cryo-SXT). Cryo-SXT allows label-free imaging of carbon- and nitrogen-containing cellular structures while preserving cells in a near-native frozen state and, as shown here, is particularly efficient in distinguishing normal and diseased-state nucleoli. The attenuation of soft X-rays into intracellular structures creates biomolecule concentration- and composition-dependent images (Loconte et al., 2022; Harkiolaki et al., 2018) from which quantitative features relevant to the material state of biomolecular condensates can be extracted. Notably, measurements of the linear absorption coefficient (LAC) provide a method to characterize nucleolar thickness and the distribution of low- and high-density regions of the nucleolus.

Cryo-SXT images and cross-sections of 3D cryo-SXT reconstruction data revealed misshaped nucleoli with an increased number of low-density foci at 24 hpi compared to noninfected cells (**Fig. 4 a and b, movie M2**). The 3D analysis of the SXT data indicated that the roughness and the total LAC value of nucleoli (**Fig. 4 c and d**) remained unchanged in infection. However, the nucleolar structure became heterogeneous with the increased appearance of infection-induced low-density foci (**Fig. 4 a** and **b**). These foci are reminiscent of the low-protein density interstices observed using TREx (**Fig. 3 a**). The number and total volume of these low-density foci increased during infection (**Fig. 4 e and f**). Sometimes, individual foci formed thin, low-density connections extending from the nucleolar rim towards the center of the nucleoli, as seen in noninfected cells (**Fig. 4 a and b**). In our TREx studies, similar structures were observed to contain intranucleolar chromatin (**Fig. 3 a and f**). Cryo-SXT analysis also revealed that the number of low-density regions was higher in infected than in noninfected cell nucleoli (**Fig. 4 g**).

**Figure 4.**
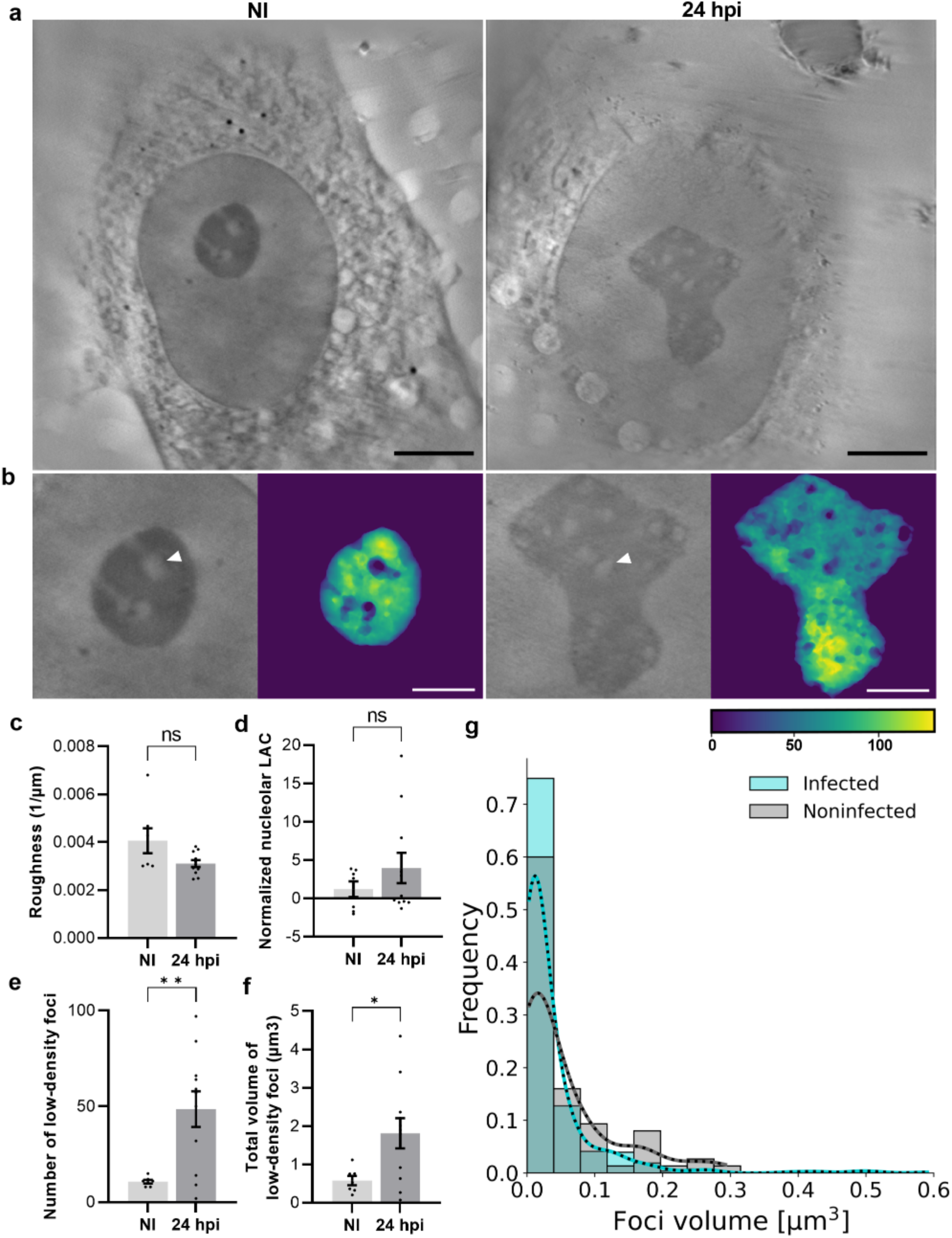
The progression of infection leads to appearance of small nucleolar foci with low density. (**a**) Cryo-soft X-ray tomography (SXT) thin slices (orthoslices) of nucleus and nucleoli in noninfected and infected NLFK cells grown on EM grids at 24 hpi. Scale bars, 5 µm. (**b**) Enlarged insets of SXT images of nucleoli and reconstruction of nucleolar thickness in noninfected and infected cells. The localization of nucleolar low-density foci and regions (filled arrowheads) are shown. The color scale indicates the thickness on the Z-axis (yellow, blue). Scale bars, 2.5 µm. See also Supplementary Movie M2. (**c**) Surface-to-volume ratio analysis of nucleolar roughness. (**d**) Nucleolar linear absorption coefficient LAC values, (**e**) the number, and (**f**) total volume of intranucleolar low-density foci. (**g**) The normalized number of various-sized nucleolar low-density foci in noninfected and infected cells (n_NI_ = 6, n_inf_ = 7). The error bars show the standard error of the mean. Statistical significance was determined using Welch’s *t*-test. The significance values shown are denoted as ** (p=0.0023), * (p=0.0114), or ns (not significant).

In conclusion, the high-resolution analysis of intranucleolar DNA and total protein content by expansion microscopy and cryo-SXT revealed the ultrastructural reorganization of nucleolar DNA and proteins, as well as the emergence of enlarged DNA-containing channels and low-density foci during infection.

### NS2 interacts with host cell proteins involved in nucleolar organization and rRNA processing

Our previous findings revealed that CPV NS2 localizes to the nucleolus during the infection and that this protein may interact with nucleolar proteins (Mattola et al., 2022b). To further investigate the extent of NS2 association with nucleolar proteins, we analyzed in greater detail our previously established NS2 interactome (Mattola et al., 2022b), now focusing on the nucleolus. Our NS2 interactome was generated by proximity-dependent biotin identification (BioID) assays (Liu et al., 2018), and mass spectrometry analyses of NS2-associated proteins. Specifically, our dataset was produced using a BirA*-tagged NS2 fusion protein expressed in Flp-In T-Rex 293 cells infected with CPV (24 hpi) and, as control, in noninfected cells.

The analysis of NS2-associated nucleolar proteins indicated that the number of proteins was slightly higher in noninfected cells (33 proteins) than in infected cells (27) (spectral count ≥ 2; Bayesian false discovery rate, BFDR, cutoff of <0.01 or <0.05) (**Fig. 5 a and b, Table T1 and T2)**. The NS2-associated proteins in noninfected cells represent putative interactions of NS2 independent of infection. In infected cells, most NS2-associated proteins with various biological functions were localized in the GC subcompartment of the nucleolus (20/27 of all spectral counts). Putative NS2 interactors were also located in the FC and DFC (six counts ≥ 2) and in the NR (seven counts ≥ 2), and some proteins showed dual localization (six) (**Fig. 5 c, Table T1 and T2)**. The highest-ranked high-confidence interactor detected by BioID was Ki-67 (MKI67), which was found in interaction with NS2 in both noninfected and infected cells (BFDR/spectral count 0.02/78.5 in infected and 0.04/76.5 in noninfected cells) (**Fig 5 a and b**, **Table T1 and T2).**

**Figure 5.**
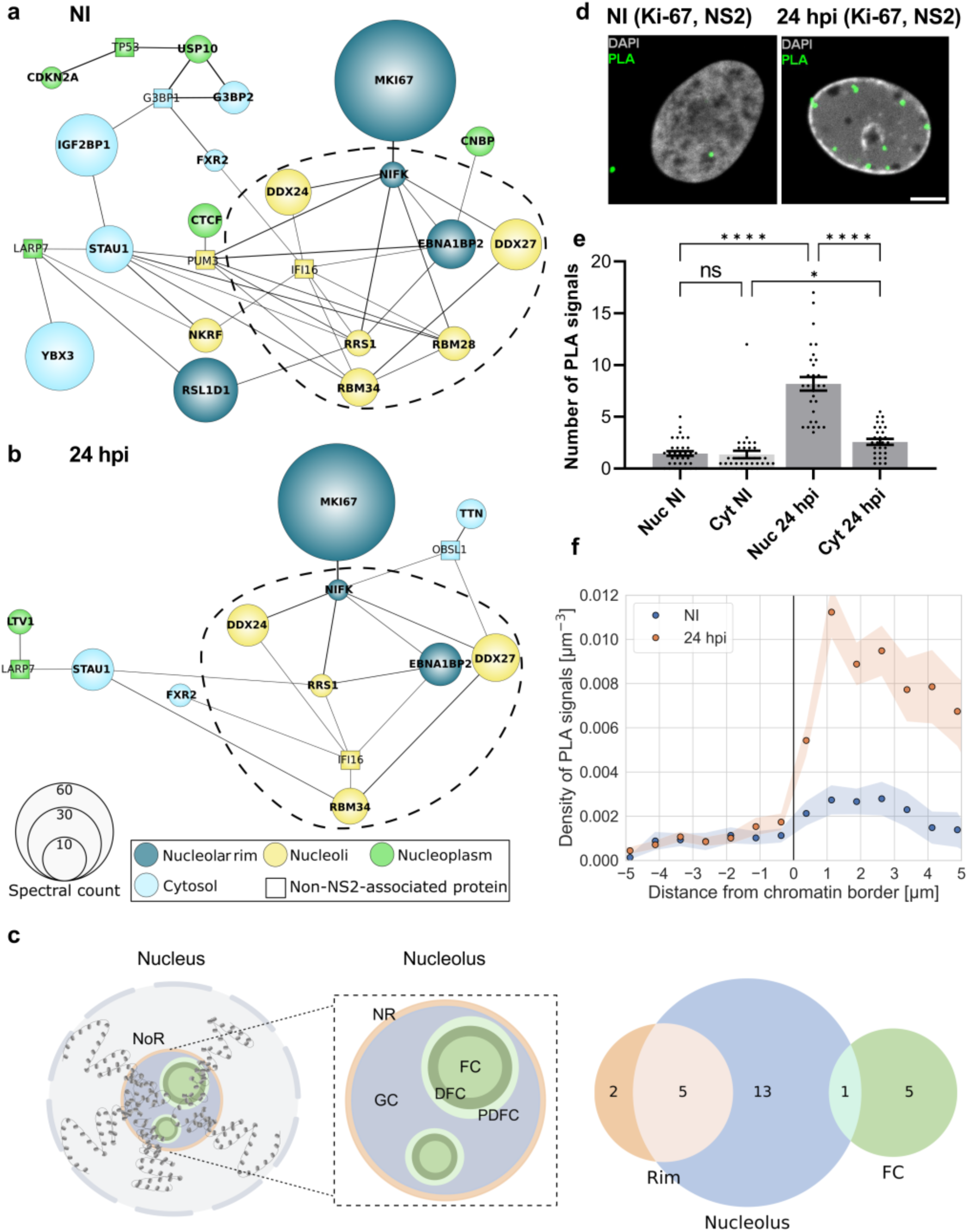
Nucleolar interactions of the viral NS2 protein. Schematic presentation of high-confidence BioID NS2 interactors related to nucleolar processes. Interactors of BirA*-tagged NS2 in (**a**) noninfected (NI) and (**b**) infected Flp-In T-REx 293 cells at 24 hpi are shown. All cells were transfected with BirA*-tagged NS2. Interactor proteins are presented by their gene names. The cellular distribution of NS2-associated nucleolar (round-shaped nodes) and nonnucleolar proteins (square-shaped nodes) in the nucleolar rim (dark blue), nucleoli (yellow), nucleoplasm (green), and cytosol (light blue) is shown. The node size correlates with interaction probability measured by spectral counts, and the black lines between nodes are proportional to the interaction in the STRING database (https://string-db.org/). Interactors are presented by their gene names. (**c**) A schematic image of the organization of the nucleolus organizer region (NoR), nucleolus, and its subcompartments: the fibrillar centre (FC), the dense fibrillar component (DFC), the periphery of dense fibrillar center (PDFC), the granular component (GC), and nucleolar rim (NR). The Venn diagram shows the nucleolar localization of NS2-associated proteins identified by BioID in infected cells at 24 hpi. The numbers of proteins localized into the whole nucleolus (blue), FC and DFC (green), nucleolar rim (orange), or dual localization (light brown, light green) are shown. (**d**) Representative confocal microscopy images showing the nuclear distribution of proximity ligation assay (PLA) foci indicating the interaction between NS2 and Ki-67 in noninfected and infected cells at 24 hpi. Scale bar, 5 μm. (**e**) The number of PLA signals in the nucleus and cytoplasm. (**f**) The density of the PLA foci as a function of increasing distance from the NE in noninfected and infected cells (n = 62 and n = 58, respectively). The error bars and the shaded areas around the data points show the standard error of the mean. Statistical significance was determined using the Games-Howell test. The significance values shown are denoted as **** (p<0.001) or * (p=0.0439), or ns (not significant).

To confirm the association between Ki-67 and NS2, we used a Proximity Ligation Assay (PLA) which generates discrete fluorescent foci at sites of interactions detectable by light microscopy only when the two antigens are within 40 nm of each other (Söderberg et al., 2006). In noninfected cells, only few foci per nucleus were detected (**Fig. 5 d**). The number of foci in infected cells increased sharply, exhibiting a highly distinctive perinuclear distribution pattern, nearly perfectly concentric. Most punctate nuclear PLA signals were localized in the nucleoplasm near the NE (**Fig. 5 d and e),** where chromatin accumulates (**Fig. 3 a**). Most of the signals were accumulated in peripheral areas, not in the center of the nucleus (**Fig. 5f)**. The negative NS2 and Ki-67 control in non-infected cells and technical probe controls (**Fig. 5 d and e, Fig. S6)** indicated that the background in the nuclear area was low.

Gene Ontology (GO) annotation analyses of biological processes indicated that 12 putative interaction partners of NS2 are involved in the organization of the nucleolus (GO:0007000, three proteins) and in rRNA processing (GO:0006364, nine proteins) (**Fig. 6 a, Table T1 and T2**). The nine proteins grouped by their involvement in rRNA processing included three DEAD-box helicases: DDX27, DDX54, and DDX56. DDX27 (BFDR <0.01) is essential for rRNA synthesis, processing, and early ribosomal assembly, and it colocalizes with FBL in the DFC region (Kellner et al., 2015; Bennett et al., 2018). We investigated if the infection affects the subcellular distribution of three NS2-associated proteins: DDX27, Epstein-Barr virus nuclear antigen 1 binding protein 2 (EBNA1BP2), and the nucleolar protein interacting with the FHA domain of Ki-67 (NIFK). In infected cells, DDX27 labeling retained the nucleolar localization but was additionally accumulated in the nucleoplasm, showing increased nuclear intensity (**Fig. 6 b and c**). EBNA1BP2 (BFDR <0.05) is an NPM1-binding protein involved in ribosomal subunit assembly (Uchihara et al., 2021; Huber et al., 2000; Kim et al., 2020). EBNA1BP2 was detected in the nucleolar area with decreased nuclear intensity in infected cells (**Fig. 6 d and e**). NIFK, the nucleolar protein interacting with the FHA domain of Ki-67 (BFDR <0.01), is an interactor of Ki-67 and NPM1 and participates in rRNA maturation colocalizing with FBL in the DFC area (Takagi et al., 2001; Sun and Kaufman, 2018; Mattola et al., 2022b; Pan et al., 2015). Upon infection, NIFK is concentrated in the nucleolar area (**Fig. 6 f and g**).

**Figure 6.**
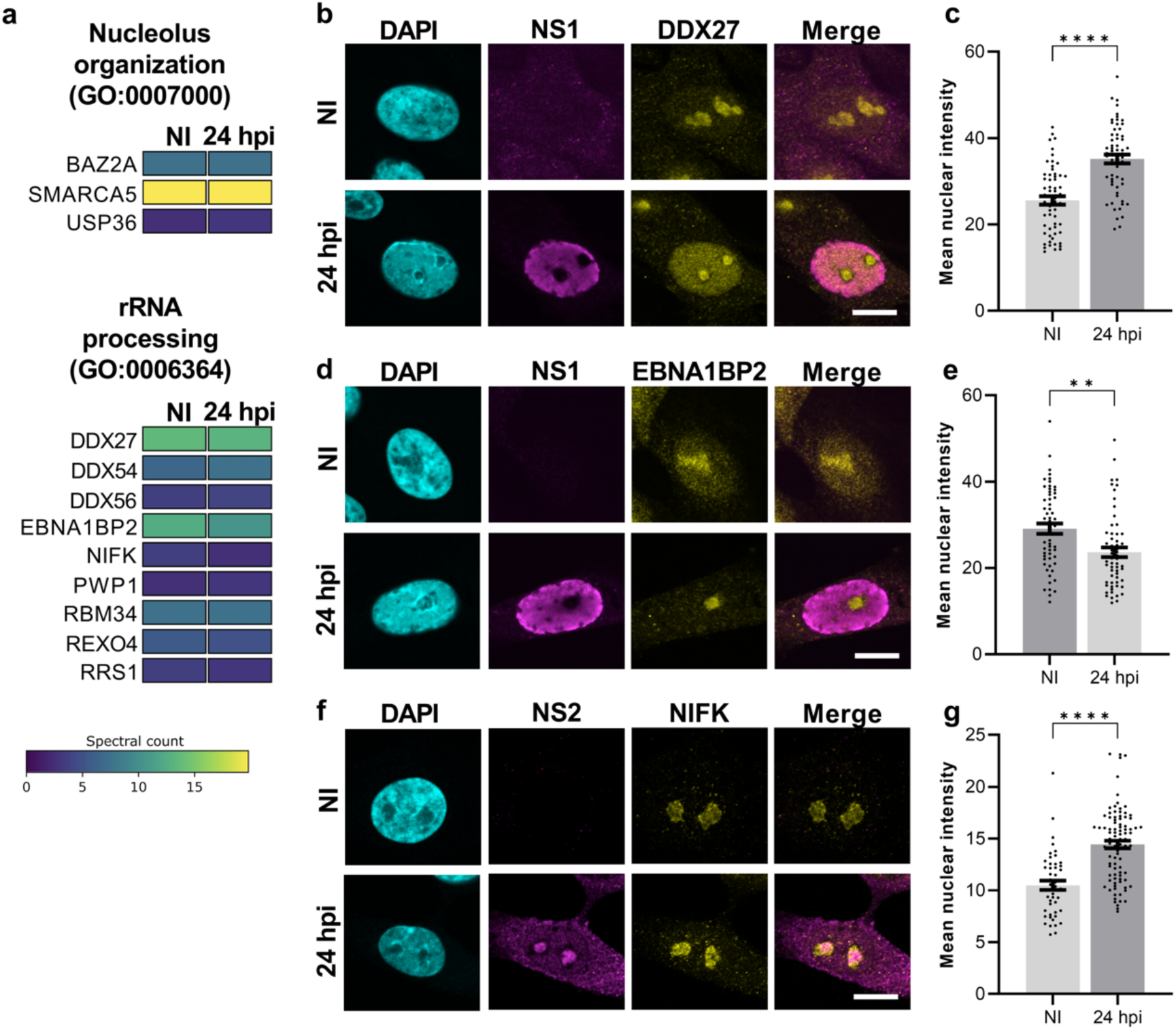
Distribution and intensity of NS2 interactor proteins are altered upon infection. (a) Identification of the viral NS2 protein nucleolar protein interactome using BioID. GO term classification for 12 high-confidence BioID NS2 interactors related to nucleolus organization and rRNA processing. Interactors of BirA*-tagged NS2 in noninfected and infected Flp-In T-REx293cells at24hpi are shown. The color bar indicates increased interaction (yellow-green) and decreased interaction (green-blue). Interactors are presented by their gene names. (b) Representative confocal images of nuclear localization of DEAD-box helicase DDX27 and (c) its mean nuclear intensity in noninfected and infected NLFK cells at 24 hpi (n=62). (d) Nuclear distribution and (e) the mean nuclear intensity EBNA1BP2 in noninfected and infected cells (n = 59). (f) Nuclear localization and (g) the mean nuclear intensity of NIFK in noninfected and infected cells (n = 47 and 92, respectively). The cells were immunolabeled with DDX27 (yellow), EBNA1BP2 (yellow), NIFK (yellow), and viral NS1 and NS2 protein (magenta) antibodies. DNA was labeled with DAPI (cyan). Scale bars, 10 µm. The error bars show the standard error of the mean. Statistical significance was determined using Welch’s t-test. The significance values shown are denoted as **** (p<0.0001) and ** (p=0.0013<0.01).

In conclusion, we detected several NS2-associated nucleolar proteins, including ribosome biogenesis factors, and showed for several of them that their association with the nucleolus is affected upon infection.

### Pre-rRNA processing is affected during CPV infection

To monitor how CPV infection may affect pre-rRNA processing, we analyzed rRNA intermediates using northern blotting in infected NLFK cells at 8, 12, and 24 hpi and, as a control, in noninfected cells (**Fig. 7**, **Fig. S7**). The pre-rRNA processing pathway has not been described in felines, so we started by designing a set of probes specific to feline pre-rRNAs, targeting all four non-coding spacers (**Fig. 7 a**). We used the rDNA of the *Felis catus* (domestic cat) genome as a reference (see Materials and methods). We modeled the feline pre-rRNA processing pathway on the well-characterized mouse one (Mullineux and Lafontaine, 2012), a closely related vertebrate.

**Fig 7.**
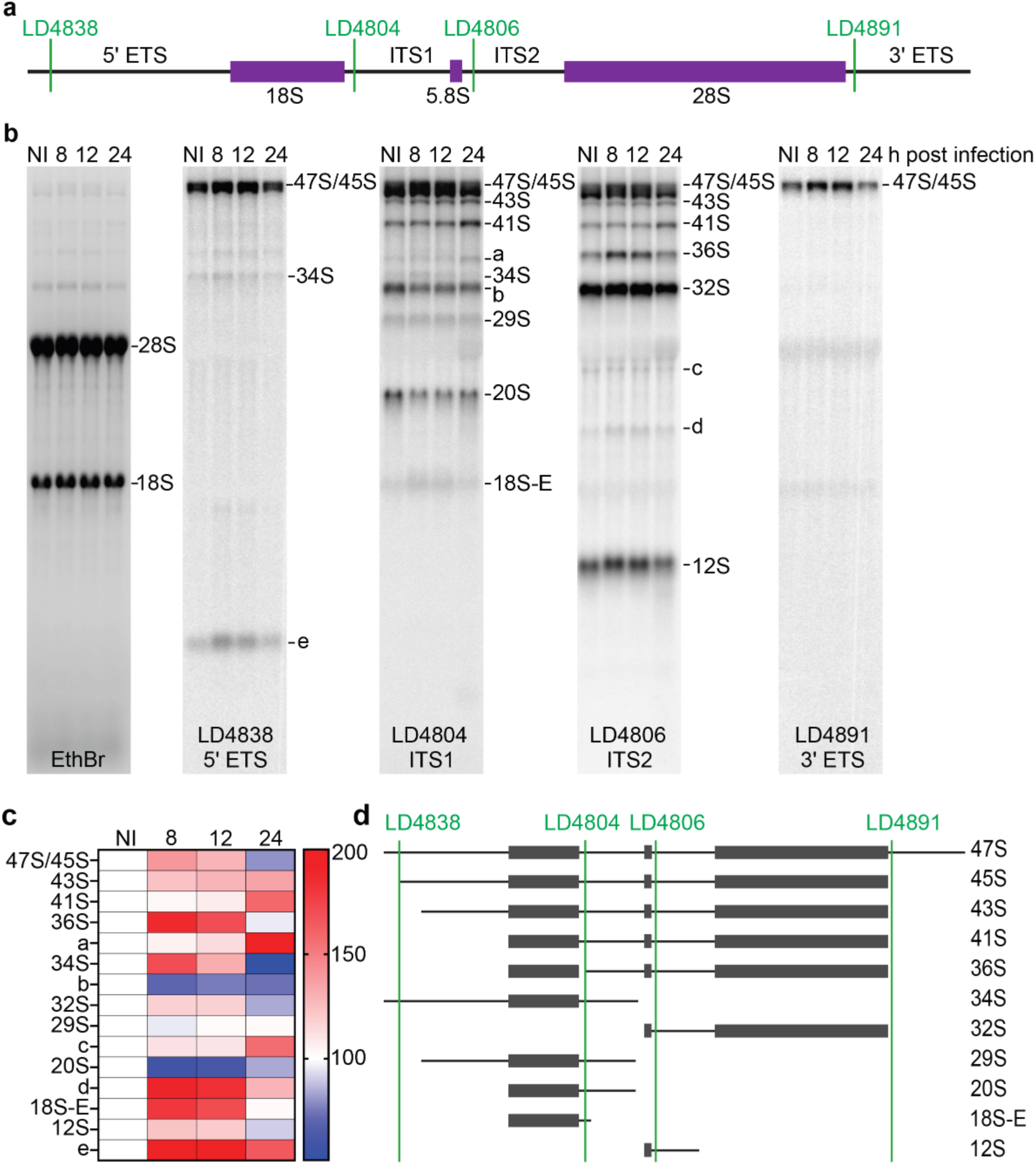
Infection alters ribosome biogenesis. (a) Schematic depiction of the feline 47S pre-rRNA and location of the probes used to detect processing intermediates. The 5’ and 3’ external transcribed spacers (ETS) and internal transcribed spacers (ITS) 1 and 2 are indicated. (b) Total RNA from NLFK cells was extracted at 8, 12, and 24 hpi. A noninfected (NI) sample was used as a control. The RNA was resolved on a denaturing high-resolution agarose gel, stained with ethidium bromide (EthBr) to reveal large mature rRNAs, or processed for northern blotting to detect precursor rRNAs. Probes for each spacer (LD4838, LD4804, LD4806, and LD4891) were designed (see Materials and Methods), revealing primary processing intermediates. Unmapped species are labeled ‘a’ to ‘e’ in order of decreasing size. (c) Phosphorimager quantification of pre-rRNA species detected. The experiment was repeated four times (n= 4, see Fig S7), and the mean values are depicted in the heatmap. A similar processing phenotype can be observed in cells 8 and 12 hpi. At 24 hpi, a different processing phenotype can be observed. (d) Putative pre-rRNA processing pathway in feline cells, based on a closely related vertebrate mouse (Inspired by Anikin and Pestov, 2022).

The most striking effects observed upon infection were: 1) a transient increase of several pre-rRNA species, observed at 8 and 12 hpi (these include the species 45S/47S, 36S, 32S, 18S-E, 12S, and species “e”), and 2) a decrease of others (notably of the 20S and “b”) (**Fig. 7 b**, and, for quantification, see heat maps on **Fig.** 7 **c**). Such effects are consistent with inhibition of pre-rRNA processing at multiple cleavages leading to a slowdown in ribosome production during viral infection (See Discussion).

These alterations of the steady-state levels of pre-rRNAs were perfectly reproducible, as shown by analysis in four independent biological replicates (**Fig. S7**). Interestingly, a return to near normal pre-rRNA levels was observed at the latest time point of infection inspected, 24 hpi (e.g., compare levels of47S/45S, 36S, or12S in the NI and 24 hpi samples). Such recovery at a late stage of infection indicates that the inhibitions caused by the presence of the virus in cells are transient in nature. Nonetheless, at this late time point post infection, the levels of41S are strikingly higher than in noninfected cells, indicating some persistent effects of infection on pre-rRNA processing (**Fig. 7 c and d, Table T3**).

In conclusion, CPV infection of cells is accompanied by a transient yet substantial inhibition of pre-rRNA processing at multiple cleavage steps, leading to an unbalanced production of pre-rRNA precursors.

## Discussion

A key feature of viruses is their ability to redirect cellular resources for their benefit, hijacking essential pathways and molecular nanomachines such as the ribosome. The nucleolus, the cell’s ribosome factory and central hub of energy consumption, is also a frequent target (Thoms et al., 2020). The nucleolus is rich in diverse catalytic activities, including folding, modification, and cleavage. Many DNA viruses, such as HSV-1, adenovirus, and AAV, are known to target the nucleolus to interfere with host cell function and to direct its proteins there to facilitate viral replication (Matthews et al., 2011; Callé et al., 2008; Salvetti and Greco, 2014; Lee et al., 2003; Wistuba et al., 1997). However, how autonomous parvovirus infection may impact nucleolar events has remained largely unknown. Our findings demonstrated that CPV infection can induce profound structural alterations of the nucleolar organization (**Figs. 1-4**) and transient alterations of pre-rRNA processing (**Fig. 7**).

### Infection impacts the distribution of nucleolar proteins and chromatin

Viral infection has been reported to lead to striking changes in the nucleolar architecture and the redistribution of nucleolar proteins (Greco, 2009; Hiscox, 2007; Atari et al., 2022; Salvetti and Greco, 2014). In HSV-1 infection, nucleolar UBF1 is relocalized from the FC to the VRC area, where it plays a role in viral replication (Stow et al., 2009). In Kaposi’s sarcoma-associated herpesvirus (KSHV), FBL is released into the nucleoplasm (Atari et al., 2022). The methyltransferase activity of FBL is also essential for cytoplasmic replication of several pathogenic viruses such as measles, mumps, and respiratory syncytial viruses (Deffrasnes et al., 2016). In HSV-1 and cytomegalovirus infections, both NCL and NPM1 become dispersed into the nucleoplasm (Callé et al., 2008; Lymberopoulos and Pearson, 2007, 2010; Bender et al., 2014; Greco et al., 2012). NPM1 is removed from the nucleoli by the endonuclease function of the HSV-1 endonuclease UL24 (Callé et al., 2008; Lymberopoulos et al., 2011). Both cellular NCL and viral UL24 are essential for the nuclear egress of HSV-1 capsids (Sagou et al., 2010; Callé et al., 2008; Lymberopoulos et al., 2011). This suggests a link between the HSV-1 infection-induced disruption of nucleolar structure integrity and viral egress.

This study focused on CPV infection and its impact on nucleolar architecture. Our findings show that the FC/DFC and NR substructures are disassembled during infection, leading to the partial release of nucleolar marker proteins into the nucleoplasm. Remarkably, during the natural disassembly of the nucleolus at the onset of mitosis in cycling cells, the FC and DFC are also the first subphases to disappear (Lafontaine et al., 2021), highlighting a parallel between nucleolar breakdown during mitosis and that induced by CPV infection. These results are consistent with our previous work, showing that CPV infection causes cell cycle arrest at the G2/M checkpoint, interfering with its release and the progression into early mitotic events (Mattola et al., 2022a).

In the nucleoplasm, the nucleolar proteins overlap with the enlarged VRC region, filling the entire nucleoplasm at the late stages of infection (**Fig. 1**). Remarkably, the PDFC and GC, markers (NCL, and NPM1, respectively) remained associated with the nucleolar area, forming protein-rich centers with increased circularity in late infection. Interestingly, NCL has earlier been shown to form complexes with the viral single-stranded genomes of MVM in vitro (Barrijal et al., 1992) and to bind AAV-2 capsids assembled in the nucleoli in vivo (Qiu and Brown, 1999; Wistuba et al., 1997; Johnson and Samulski, 2009). Our findings show that the inner core of the nucleolus and the NR lose integrity upon CPV infection, whereas the peripheral layers remain relatively intact. This suggests that in contrast to HSV-1 infection, CPV infection induces only a partial disruption of nucleolar integrity. However, the increased circularity of the nucleolar condensates containing NCL, NPM1, and pre-rRNAs indicates an alteration of surface tension of these foci and a change in phase behavior (**Fig. 2**). These findings suggest that parvovirus infection changes nucleolar organization with consequences on ribosome biogenesis.

The probing of nucleolar ultrastructure by TREx and cryo-SXT revealed that the progression of infection leads to extensive structural remodeling of nucleoli, including condensation of chromatin around the nucleoli. The observation using TREx microscopy showed that the infection also resulted in the emergence of enlarged nucleolar low-protein density interstices and their vacuolar cavities (**Fig. 3 and 4**). Similar structural changes have been observed in stressed cells after vitamin C and K3 (Apatone) and actinomycin D treatments (Jamison et al., 2010; Tchelidze et al., 2017). The emergence of interconnected channels forming a network is compatible with the notion that the nucleolus has been described as a percolated biomolecular condensate. It is interesting to observe that the size and distribution of the internal cavities seem to change drastically, becoming larger upon viral infection as the nucleolus is remodeled. The virus-induced compaction of intranucleolar DNA could limit chromatin accessibility to transcription factors and thereby downregulate the level of rRNA gene transcription. The redistribution of nucleolar UBF1, a regulator of transcription of rDNA (Sanij and Hannan, 2009; Sanij et al., 2008), along with chromatin reorganization, may reducer RNA transcription. However, our findings show that the total intensity of nucleolar rRNA and the production of primary rRNA transcripts remain relatively unchanged in infection. This may suggest that the partial compaction of intranucleolar chromatin in infected cells does not necessarily lead to significant interference with rRNA transcription. However, the infection may impact the pre-rRNA processing pathway.

### CPV NS2 interacts with nucleolar proteins

During the infection of various viruses, several proteins localize to the nucleolus, where they interact with nucleolar proteins (Salvetti and Greco, 2014; Iarovaia et al., 2021). These interactions lead to the virus-induced disruption of nucleolar function and recruitment of cellular proteins to aid viral replication. During henipavirus infection (RNA virus family *Paramyxoviridae*), the viral matrix proteins interact with Treacle, a nucleolar protein that regulates rDNA transcription (Rawlinson et al., 2018). The acquisition of Treacle during evolution is suggested to coincide with the emergence of the FC subcompartment of the nucleolus (Lafontaine, 2023b; Jaberi-Lashkari et al., 2023). In HSV-1 infection, NPM1 translocation outside ofthe nucleolus is assisted by the viral protein UL24, and that of NCL by viral ICP4 and US11 (Callé et al., 2008; Lymberopoulos et al., 2011; Greco et al., 2012). The Epstein-Barr virus protein EBNA11 recruits NPM1 for transcriptional activation of its latency genes (Malik-Soni and Frappier, 2014).

Detailed analysis ofCPVNS2 interactome (Mattola et al., 2022b) suggested that NS2 interacts with several proteins of the nucleolar inner core, peripheral layers, and NR(**Fig. 5**, and Mattola et al. 2022). These putative nucleolar interactors ofNS2 are involved in nucleolar organization and rRNA processing. A major NS2 interactor is the chromosome surfactant protein Ki-67, a protein involved in the organization of heterochromatin in actively dividing mammalian cells (Sobecki et al., 2016; van Schaik et al., 2022; Booth et al., 2014). It also has a role in the nucleolar assembly and organization of perinucleolar chromatin at the NR (van Schaik et al., 2022; Remnant et al., 2021). NS2 was also associated with cellular proteins involved in ribosome biogenesis and rRNA processing, whose subcellular distribution was modified upon CPV infection (**Fig. 6**). The interaction ofNS2 with proteins regulating nucleolar organization suggests that NS2 has a role in the profound alteration of nucleolar morphology and function.

### CPV infection impacts pre-rRNA processing

Nucleolar structure alterations during CPV infection impacted the function of the nucleolus (**Fig. 7**). There was a transient change in the steady-state levels of most pre-rRNA precursors, which is compatible with the inhibition of cleavage at multiple processing steps. Interestingly, the observed effects on processing appeared transient at 8 and 12 hpi but not later. Remarkably, the steady-state levels of most pre-rRNA returned to near normal levels at 24 hpi at a time of infection when the nucleolar structure was still profoundly impacted.

#### Why might it be interesting for the virus to repress host cell ribosome biogenesis during infection?

Firstly, it is essential to consider that ribosomes are long-lived (i.e., several days) ribonucleoprotein particles that are robustly packaged during assembly. This implies that pre-existing ribosomes remain available to produce the viral components throughout infection. Secondly, ribosome biogenesis is an energy-intensive process that engages most cellular activities, including all three RNA polymerases and hundreds of factors involved in RNA cleavage, modification, folding, and transport – some are also essential for other cellular substrates. Additionally, ribosome biogenesis spans multiple cellular subcompartments: it begins in the nucleolus, continues in the nucleoplasm, and is finalized in the cytoplasm. Thus, it is reasonable to assume that suppressing ribosome biogenesis, at least temporarily, is beneficial for the virus, as it reduces the host cell’s energy expenditure and redirects resources toward processes that favor viral replication.

In conclusion, our work demonstrates that autonomous parvoviruses, such as CPV, not only remodel the structural organization of the nucleolus but also modulate its function. This study provides novel insights into the functional and physical interactions between autonomous parvoviruses and the nucleolus.

## Materials and methods

### Cell lines, plasmids, and viruses

Norden laboratory feline kidney (NLFK) cells were maintained as previously described (Mattola et al., 2022b). Infectious plasmid clone pBI265 (Parrish, 1991) was used to generate CPV-2. Cells were transfected using JetOptimus transfection reagent (Polyplus-transfection, Illkirch, France) as described in the manufacturer’s instructions. Virus stocks were produced in NLFK cells and concentrated using an Amicon® Stirred Cell ultrafiltration system (Merck KGaA, Darmstadt, Germany).

### Antibodies

The following primary antibodies against viral antigens used in confocal microscopy analyses were generous gifts: mouse anti-NS1 (Dr. Caroline Astell, University of British Columbia, Vancouver, Canada) (Yeung et al., 1991). mouse anti-NS2 and rabbit anti-VP2 (Dr. Colin Parrish, Cornell University, Ithaca, NY, USA) (Wang et al., 1998; Weichert et al., 1998). Commercial primary antibodies against cellular proteins were used according to the manufacturer’s instructions: rabbit UBF1 (ab244287, Abcam, Cambridge, UK), rabbit FBL (ab166630, Abcam), rabbit NCL (ab22758, Abcam), mouse NPM1 (conjugated to Alexa647, MA3-25200-A647, Invitrogen, Life Technologies, Waltham, MA), rabbit Ki-67 (ab15580, Abcam), mouse rRNA (MA1-16628, Invitrogen) (Garden et al., 1995), rabbit DDX27 (17087-1-AP, ProteinTech Europe, Manchester, UK), rabbit EBNA1BP2 (PA5-97189, Invitrogen), rabbit NIFK (HPA 035735, Sigma, Merck KGaA). The primary antibodies were followed by goat anti-mouse or anti-rabbit Alexa448, Alexa546, and Alexa647 conjugated secondary antibodies (Thermo Fisher Scientific, Waltham, MA, USA).

### Immunolabelling and confocal microscopy

For microscopy sample preparation, NLFK cells were cultured on glass coverslips at 37°C with 5 % CO^2^. Cells were infected with CPV and fixed at 8, 12, or 24 hpi in 4% paraformaldehyde for 12 minutes. Following fixation, cells were permeabilized with 0.1% Triton X-100 in phosphate-buffered saline supplemented with 0.1% bovine serum albumin. After primary antibody immunolabeling, cells were labeled with mouse or anti-rabbit secondary fluorescent antibodies. DNA was stained with Pro-Long Diamond anti-fade media with DAPI (Thermo Fisher Scientific).

The microscopy of immunolabeled samples was performed with Leica TCS SP8 FALCON (Leica microsystems, Mannheim, Germany) and Nikon A1R (Nikon, Tokyo, Japan) laser scanning confocal microscopes. The parameters for images acquired with Leica TCS SP8 FALCON were as follows: DAPI was excited with a 405 nm diode laser, and the fluorescence was collected between 415-495 nm. Alexa488 and Alexa546 were excited with 499 nm and 557 nm wavelengths of pulsed white light laser (80 MHz). The emission detection range was adjusted to 505-561 nm for Alexa488 and 568-648 nm for Alexa546. Microscopy images with Nikon were acquired by exciting DAPI with a 405 nm diode laser and collecting with a 450/50 nm band-pass filter. Alexa488 was excited with a 488 nm argon laser, and the fluorescence was collected with a 515/30 nm band-pass filter. Alexa 647 was excited with a 640 nm diode laser and detected with a 660 long-pass filter. The image resolution was 512 x 512 or 1024 x 1024 pixels, with a pixel size between 30-60 nm in the x- and y-directions and 180-300 nm in the z-direction. A fixed optical configuration was used to acquire all images of the same sample set.

The nuclei were segmented from the DAPI stain using Otsu’s or minimum cross entropy method (Otsu, 1979; Li and Lee, 1993). Total fluorescence intensities in the nucleus and nucleoli were calculated by summing the pixel intensity values within the segmented regions. The foci of nucleolar proteins were segmented by adjusting the intensity threshold to 33%, 45%, and 55% of the intensity range in each nucleus for UBF1, fibrillarin, and nucleolin, respectively. rRNA signal was segmented using Otsu’s algorithm. The intensity distributions of labels with respect to the nuclear or nucleolar border were determined by calculating the shortest distance of each pixel to the border and calculating mean intensity as a function of the distance using 2-pixel wide distance bins for averaging. Due to the flat shape of fixed cells, the confocal section with the largest nuclear area was selected for the analyses of intensity with respect to the nuclear border, and 2D Euclidean distance was used as a metric. Since there can be many nucleoli in one cell and the nucleoli can lie on various confocal planes, the whole 3D stack was used in the analyses for the nucleolar border, and 3D Euclidean distance was used as a metric. Correlations between DNA staining and antibody labels were determined by using Pearson correlation coefficients in the nuclear region. For all the analyses, the given statistics were calculated over cells. The circularity of the single planes of segmented rRNA antibody-labeled nucleoli was determined with Fiji (Schindelin et al., 2012). Circularity determination in Fiji is based on the equation 4pi(area/perimeter^Λ^2), where a perfect circle is indicated by the value 1.0.

### Ten-fold robust Expansion Microscopy and visualization

NLFK cells were grown on coverslips as previously. Noninfected and infected cells were fixed at 24 hpi in 4% PFA (Electron Microscopy Sciences, SKU 15710, diluted to 4% with sterile PBS) for 12 min at room temperature, after which the cells were permeabilized with 0.5% Triton X-100 in PBS supplemented with 0.5% bovine serum albumin (BSA). The permeabilized cells were immunolabelled with mouse anti-NS1 and rabbit anti-fibrillarin primary antibodies, and anti-mouse Alexa546 and anti-rabbit Alexa448 secondary antibodies. Primary and secondary antibodies were used at 3-fold and 4-fold higher concentrations, respectively, compared to standard immunolabeling.

Ten-fold Robust Expansion Microscopy was performed according to the protocol described earlier (Damstra et al., 2022). Briefly, immunolabeled samples on glass coverslips were anchored using 0.1 mg/ml Acrolyl-X (Invitrogen, Catalog number A20770) overnight at + 4°C. Gelation was performed with a gel containing 14.2% acrylamide (Sigma-Aldrich, product number A9099), 10.1% sodium acrylate (AmBeed, catalog number A107105), 0.005% bis-acrylamide (Sigma-Aldrich, product number M1533), 0.15% ammonium persulfate (Sigma-Aldrich, A3678), and 0.15% TEMED (Sigma-Aldrich, product number T22500) in PBS for 1 hour at 37 °C. After gelation, the samples were denatured in 0.2 M NaCl, 50 mM Tris (pH 6.8), and 5.76% SDS for 1.5 hours at 78°C and washed twice with PBS. Gels were stained for the total protein with 20 μg/ml of ATTO 647N NHS (Atto-tec, Siegen Germany) dye in sodium bicarbonate buffer (Sigma) for 1 hour at RT. DAPI (Roche) diluted 1:6000 in PBS was used to stain DNA for 60 minutes before the expansion. Gels were expanded in distilled H_2_O, changing the water until no further expansion was observed. The achieved expansion factor (approximately 7.5) was calculated by comparing the gel size after the gelation and after the expansion and back-calculated into image scale bars.

Images of expansion samples were acquired with a Leica SP8 confocal microscope using the 63x water immersion objective (1.2 NA). DAPI was excited with a 405 nm laser and detected with a PMT detector at 412-480 nm. Alexa 488 was excited with a 499 nm laser and detected at 506-552 nm using a Leica HyD SMD detector. Alexa 546 was excited with a 557 nm laser and detected at 566-611 nm using a Leica HyD SMD detector. The total protein stain was excited with a 645 nm laser and detected at 652-800 nm using a Leica HyD SMD detector. The pixel size was 90 nm in the x- and y-directions and 356 nm in the z-direction. The images were first denoised using a Gaussian filter with a radius of three pixels to analyze the DNA and total protein labels in the nucleoli. Then, the nucleoli were segmented by setting an intensity threshold for the total protein channel using Otsu’s algorithm (Otsu, 1979). In segmenting cells with contiguous nucleoli and strong total protein signal at the nuclear envelope, the total protein images were manually edited to remove the continuity and prevent the segmentation of other regions besides the nucleoli. The coefficient of variation (the standard deviation divided by the mean) for the DNA and total protein labels and the Pearson correlation coefficient were calculated for the pixels belonging to the segmented nucleolar region. The intensity of the labels for the nucleolar border was analyzed by calculating the minimum distance of each pixel to this border, grouping the pixels based on their distance to the border, and calculating the mean intensity of each group. Surface renderings of the noninfected and infected nucleoli were created with Dragonfly 3D World v2024.1.

### Cryo-SXT of adherent cells on grids

NLFK cells were grown and infected with CPV for 24 hours at 37**°**C on Au grids (Quantifoil-coated Au-TEM finder grids, R2/2, G200F1, Gilder, Jena, GER). Noninfected and infected cells were fixed with 4% PFA for 12 minutes at room temperature and labeled with 1µg/ml Hoechst 33342. The infected cells were identified by an infection-induced chromatin marginalization using a widefield microscope (Leica DM IL LED, Leica Microsystems GmbH, Germany) with HC FL PLAN 40X/0.65NA air objective or a confocal microscope (Leica SP 8, Leica Microsystems GmbH, Germany) with HC PL APO 63x/1.2NA water immersion objective. The grids were plunged into liquid ethane for vitrification (Leica EM GP2, Leica Microsystems GmbH, Germany) and kept at -196°C until imaging. Using a compact soft X-ray microscope, the pre-vitrification-located infected cells were re-located by a low magnification SXT imaging, and a built-in cryo-fluorescence microscopy from the vitrified grids, and tomographic tilt-series of infected cells were collected (SXT100, Sirius XT, Dublin, IRE, https://siriusxt.com/) (O’Connor et al., 2024; Fahy et al., 2021). The tilt series were acquired assets of X-ray projections covering a 120° tilt range with one-degree increments. The exposure per projection ranged from 15 to 60 seconds, depending on the sample ice thickness and tilt angle. The exposure time was increased to 120 seconds at high tilt angles for samples with particularly thick ice. The samples were maintained at temperatures of - 1 63°C or lower throughout all experimental stages. The tomographic data alignment and data analysis were done as described in detail (Leclerc et al., 2024).

### BioID analyses

BioID of CPV NS2 protein was performed as previously described (Mattola et al., 2022b). Briefly, a biotin protein ligase was fused to NS2 to allow for the detection of proximal and interacting proteins by biotin tags. The high-confidence NS2 interactions identified among the biotinylated proteins were further screened for nucleolar interactors. Data was analyzed with SAINTexpress (Teo et al., 2014), and filtered using a protein-level Bayesian FDR of 0.05 as the cutoff. Controls included four different GFP-BirA* constructs, including NLS-GFP. Further filtering was done by integrating control experiment data from the CRAPome contaminant repository (Mellacheruvu et al., 2013). The high-confidence interactor set was used for Protein ANalysis THrough Evolutionary Relationships (PANTHER) classification system (http://www.pantherdb.org/) to create GO biological process annotation chart of NS2 interactors (**Table T1**).

### Proximity ligation assay (PLA)

The association ofNS2 with the key interactor Ki-67 during infection was verified with PLA. NLFK cells were grown on coverslips and infected with CPV. Infected and noninfected samples were fixed at 24 hpi in 4% formaldehyde for 12 min and permeabilized using 0.1% Triton X-100 in PBS. NS2 and Ki-67 were identified with mouse anti-NS2 and rabbit anti-Ki-67 antibodies, respectively, and the assay was performed using the Duolink In Situ Orange Mouse/Rabbit kit (Merck KGaA) following the manufacturer’s protocol. Technical controls were prepared from infected samples by omitting the anti-NS2 antibody or by omitting the PLA probes. Samples were embedded into a non-hardening mounting medium containing DAPI and imaged with a Nikon A1R confocal microscope with Plan Apo VC 60x oil immersion objective (NA 1.4). The human-in-the-loop machine-learning-based characterization of DAPI-labeled chromatin distribution verified the infection in PLA-labeled cells. The machine learning model was built, trained, and tested using Keras 1.0 with TensorFlow 2.0 backend using Python 3.9 in the Google Colab pro-environment. The deep learning code and model developed in this study can be found at https://github.com/leclercsimon74/Infection_Prediction_CPV. DAPI was excited with a 405 nm diode laser, and the fluorescence was collected with a 450/50 nm band-pass filter. PLA signals were excited with a 561 nm sapphire laser, and the fluorescence was collected with a 595/50 nm band-pass filter. Single cells were imaged as stacks (resolution 512 x 512 pixels, pixel size in xy 60 nm, z sampling distance 200 nm).

DAPI signal was used to segment the nuclei from image stacks by automatic minimum cross entropy segmentation (Li and Lee, 1993) and filling any holes within the segmented regions. The PLA signals were segmented using the maximum entropy algorithm (Kapur et al., 1985), and their geometric centers were determined. The number of PLA signals with their geometric center located within the segmented nucleus was counted.

### Quantitative northern blotting analysis of pre-rRNA

Total RNA was extracted from noninfected and infected NLFK cells at 8, 12, and 24 hpi with TRI reagent solution (Thermofisher), according to the manufacturer’s instructions. To analyze high-molecular-weight RNAs, 5 μg total RNA was resolved in a 1.2% formaldehyde-agarose denaturing gel, followed by electrophoresis at 65 V. Agarose gels were transferred overnight by capillarity onto Hybond-N+ membranes (Cytiva) in 10x saline sodium citrate (SSC). Membranes were prehybridized at 65°C for 1 h in a solution containing 50% formamide, 5x SSPE, 5x Denhardt’s solution, 1% (wt/vol) SDS, and 200 mg/ml fish sperm DNA (Roche). Hybridization with specific ^32^P-labeled oligonucleotide probes was carried out for 1 h at 65°C and overnight at 37°C. Probe sequences are listed in Table S1. The membranes were washed and exposed to Fuji imaging plates (Fujifilm). Signals were captured with a Phosphorimager (FLA-7000, Fujifilm) and quantified with Multi Gauge Software (Fujifilm, v 3.1).

### Probe sequence Table

The probes LD4838, LD4804, LD4806, and LD4891 were designed using the Felis catus_Fca126_mat1.0 genome (NCBI RefSeq assembly: GCF_018350175.1). Their sequence is given in **Table S1**.

**Table S1.**
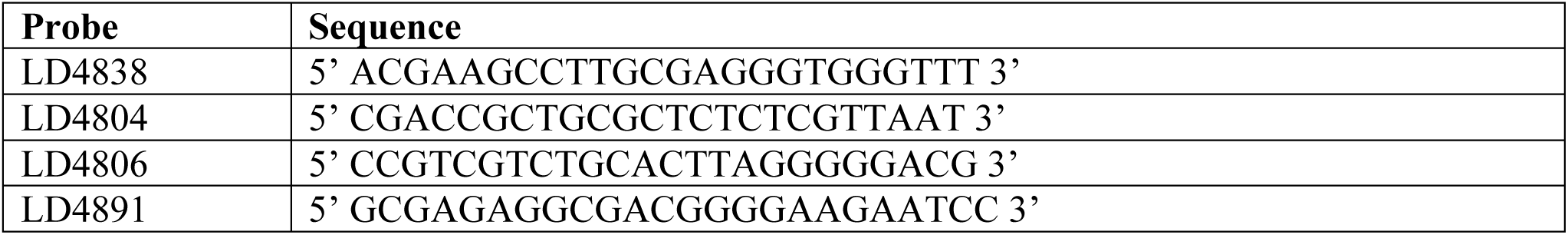
Probes used in this work.

Quantification of pre-rRNA processing intermediates (Supplementary Table 3)

## Data availability

All relevant data and supporting information files are in the manuscript.

## Contributions

**Conceptualization:** Vesa Aho, Salla Mattola, Denis Lafontaine, Maija Vihinen-Ranta

**Data curation:** Salla Mattola, Moona Huttunen, Satu Hakanen, Aynur Soenmez, Sergey Kapishnikov, Visa Ruokolainen, Denis Lafontaine,

**Formal analysis:** Simon Leclerc, Kari Salokas, Vesa Aho

**Funding Acquisition:** Colin R Parrish, Lafontaine, Maija Vihinen-Ranta

**Investigation:** Salla Mattola, Moona Huttunen, Satu Hakanen, Aynur Soenmez, Alessandro Zannotti, Sergey Kapishnikov, Visa Ruokolainen, Kari Salokas

**Methodology:** Salla Mattola, Simon Leclerc, Kari Salokas, Markku Varjosalo, Kenneth Fahy, Denis Lafontaine

**SXT data acquisition:** Alessandro Zannotti

**Software:** Simon Leclerc, Vesa Aho

**Resources:** Markku Varjosalo, Colin R Parrish, Kenneth Fahy, Denis Lafontaine, Maija Vihinen-Ranta

**Project Administration:** Maija Vihinen-Ranta

**Visualization:** Salla Mattola, Moona Huttunen, Satu Hakanen

**Writing – Original Draft Preparation:** Denis Lafontaine, Maija Vihinen-Ranta

**Writing – Review & Editing:** Moona Huttunen, Satu Hakanen, Leena Latonen, Vesa Aho, Salla Mattola, Denis Lafontaine, Maija Vihinen-Ranta

The experiments were done using commercially available cell lines without ethical challenges.

## Electronic supplementary material

Supplementary material includes methods, supplementary figures, tables, and movies.

## Acknowledgments

We thank the staff of the Biocenter Finland and Proteomics unit (Helsinki Institute of Life Science, HiLIFE; Institute of Biotechnology Proteomics Unit; Helsinki; Finland) for providing the Bio-ID analysis.

---

**Figure S1.**
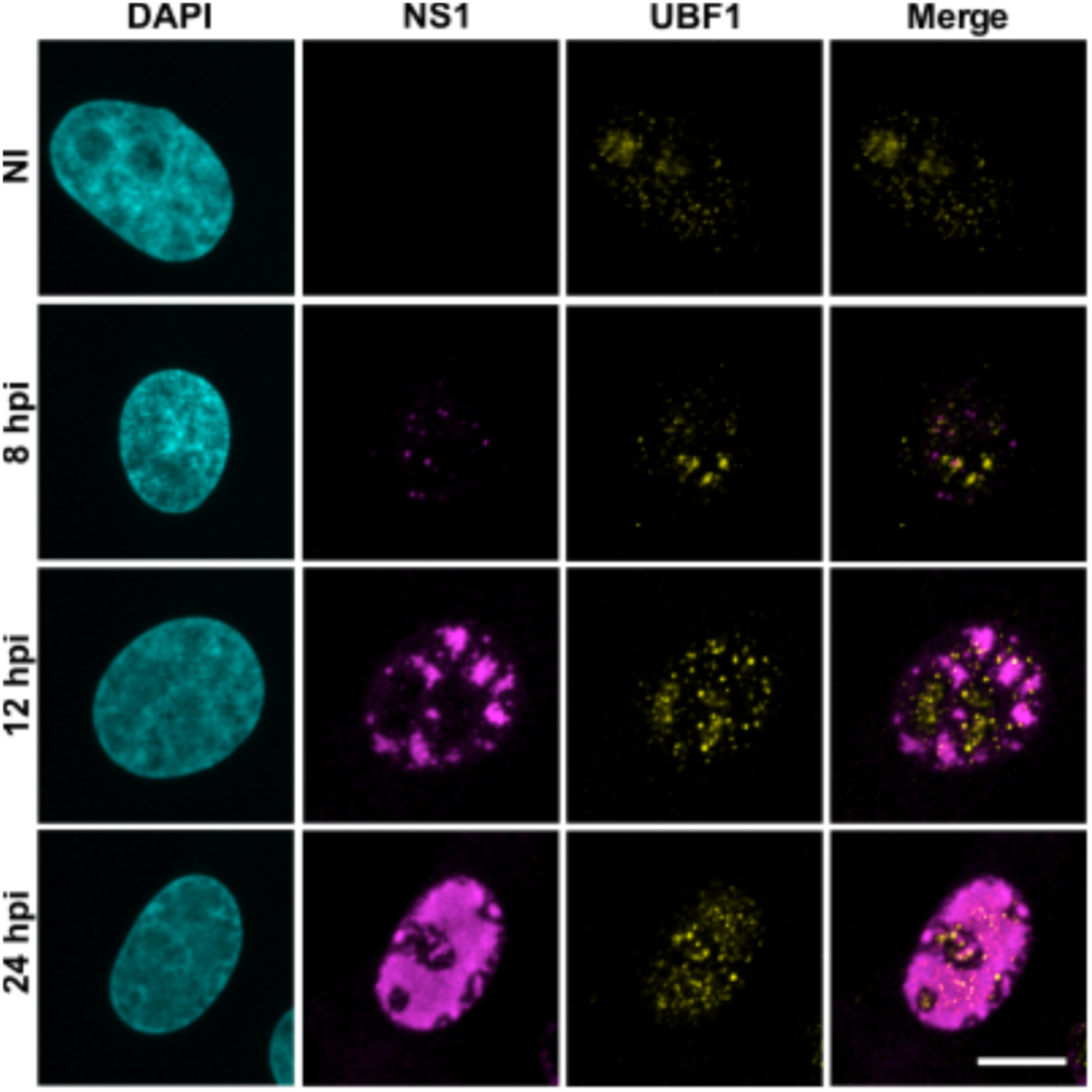
The inner core of the nucleolus is disassembled upon CPV infection, leading to redistribution of UBF1 in the nucleoplasm. Representative confocal maximum intensity projections showing FC marker protein, upstream-binding factor 1 (UBF1), localization in noninfected (NI) and infected NLFK cells at 8, 12, and 24 hpi. The cells were immunolabeled with UBF1 (yellow) and the viral NS1 protein (magenta) antibodies. DNA was labeled with DAPI (cyan). In the NS1 channel, individual VRC (white arrow, 8 hpi) progressively merge into larger structures (12 hpi) to fill in the entire nucleoplasm (24 hpi) as the infection proceeds. Scale bar, 10 µm.

**Figure S2.**
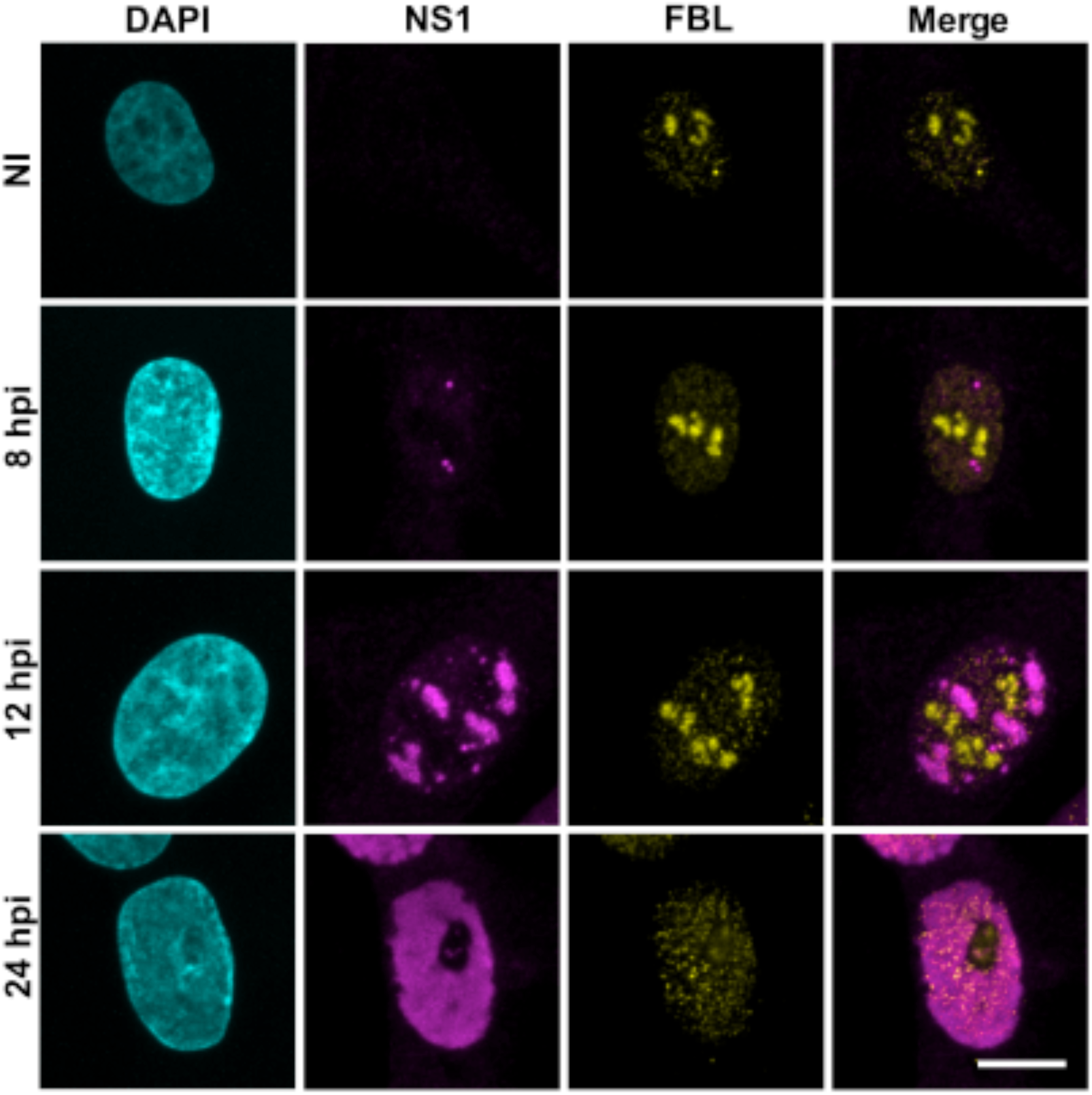
The inner core of the nucleolus is disassembled upon infection, leading to redistribution of fibrillarin in the nucleoplasm. Representative confocal maximum intensity projections showing the nucleolar localization of DFC marker protein, fibrillarin (FBL), in noninfected and infected NLFK cells at 8, 12, and 24 hpi. The cells were immunolabeled with fibrillarin (FBL, yellow) and the viral NS1 protein (magenta) antibodies. DNA was labeled with DAPI (cyan). Scale bar, 10 µm.

**Figure S3.**
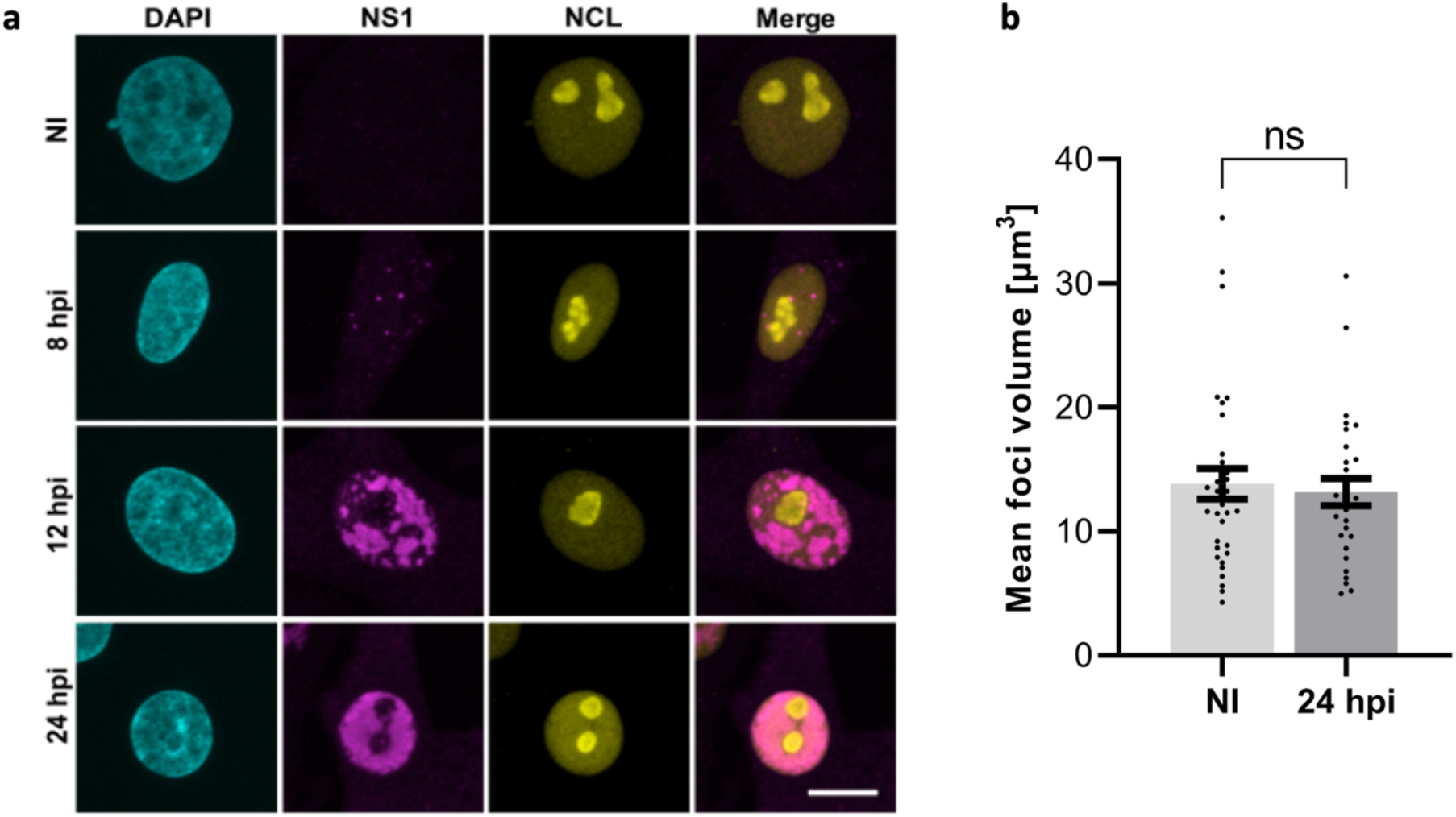
The periphery of the nucleolus is not disassembled upon infection. (**a**) Representative confocal maximum intensity projections showing the localization of peripheral PDFC marker protein, nucleolin (NCL), in noninfected and infected NLFK cells at 8, 12, and 24 hpi. The cells were immunolabeled with nucleolin (NCL, yellow) and the viral NS1 protein (a marker of viral replication compartment region, magenta) antibodies. DNA was labeled with DAPI (cyan). Scale bar, 10 µm. (**b**) The volume of NCL foci in noninfected and infected cells at 24 hpi (n = 34 and 29, respectively). The error bars show the standard error of the mean. No statistically significant difference was observed by Welch’s *t*-test (p=0.6927)

**Figure S4.**
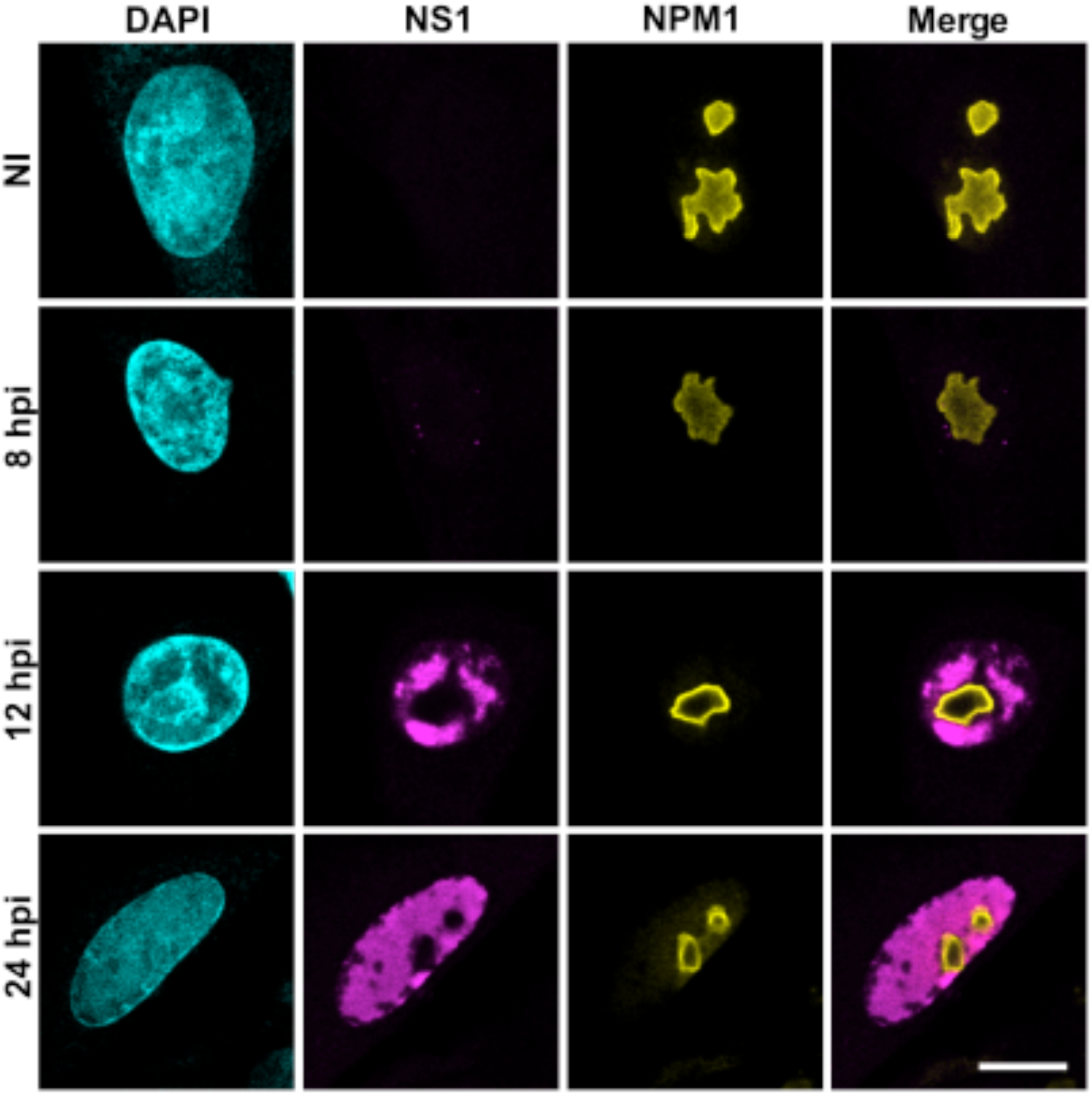
The periphery of the nucleolus is not disassembled upon infection. Representative confocal images of cellular localization of GC marker protein, nucleophosmin 1 (NPM1), in noninfected and infected NLFK cells at 8, 12, and 24 hpi. The cells were immunolabeled with NPM1 (yellow) and the viral NS1 protein (magenta) antibodies. DNA was labeled with DAPI (cyan). Scale bar, 10 µm.

**Figure S5.**
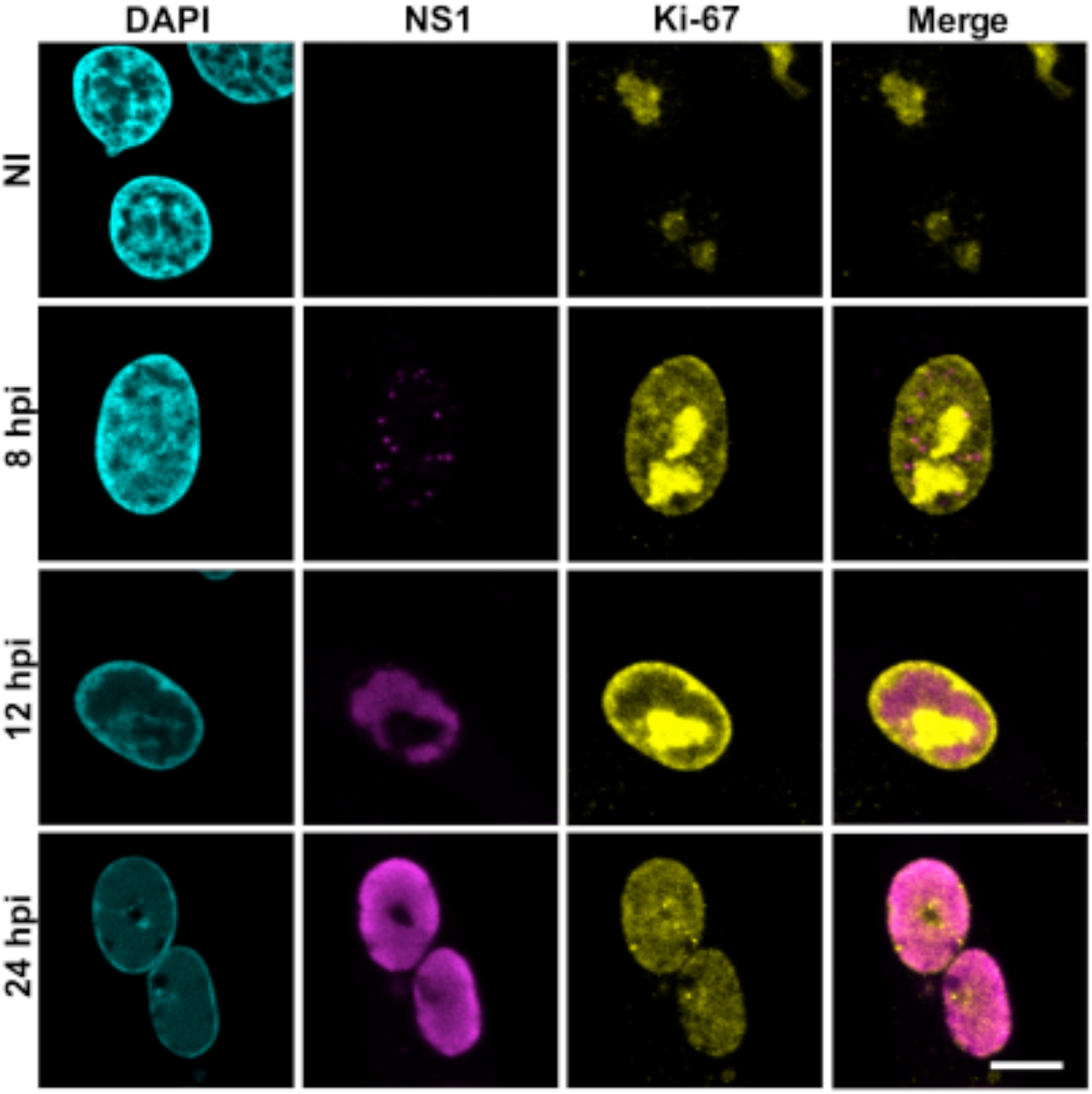
The nucleolar rim protein Ki-67 looses its association with the nucleolus at late infection. Representative confocal images of cellular localization of NR marker protein, Ki-67, in noninfected and infected NLFK cells at 8, 12, and 24 hpi. Scale bar, 10 µm. The cells were immunolabeled with Ki-67 (yellow) and the viral NS1 protein (magenta) antibodies. DNA was labeled with DAPI (cyan). Scale bar, 10 µm. Note that at 12 hpi, Ki-67 is strikingly redistributed at the nuclear envelope (see text for details).

**Figure S6.**
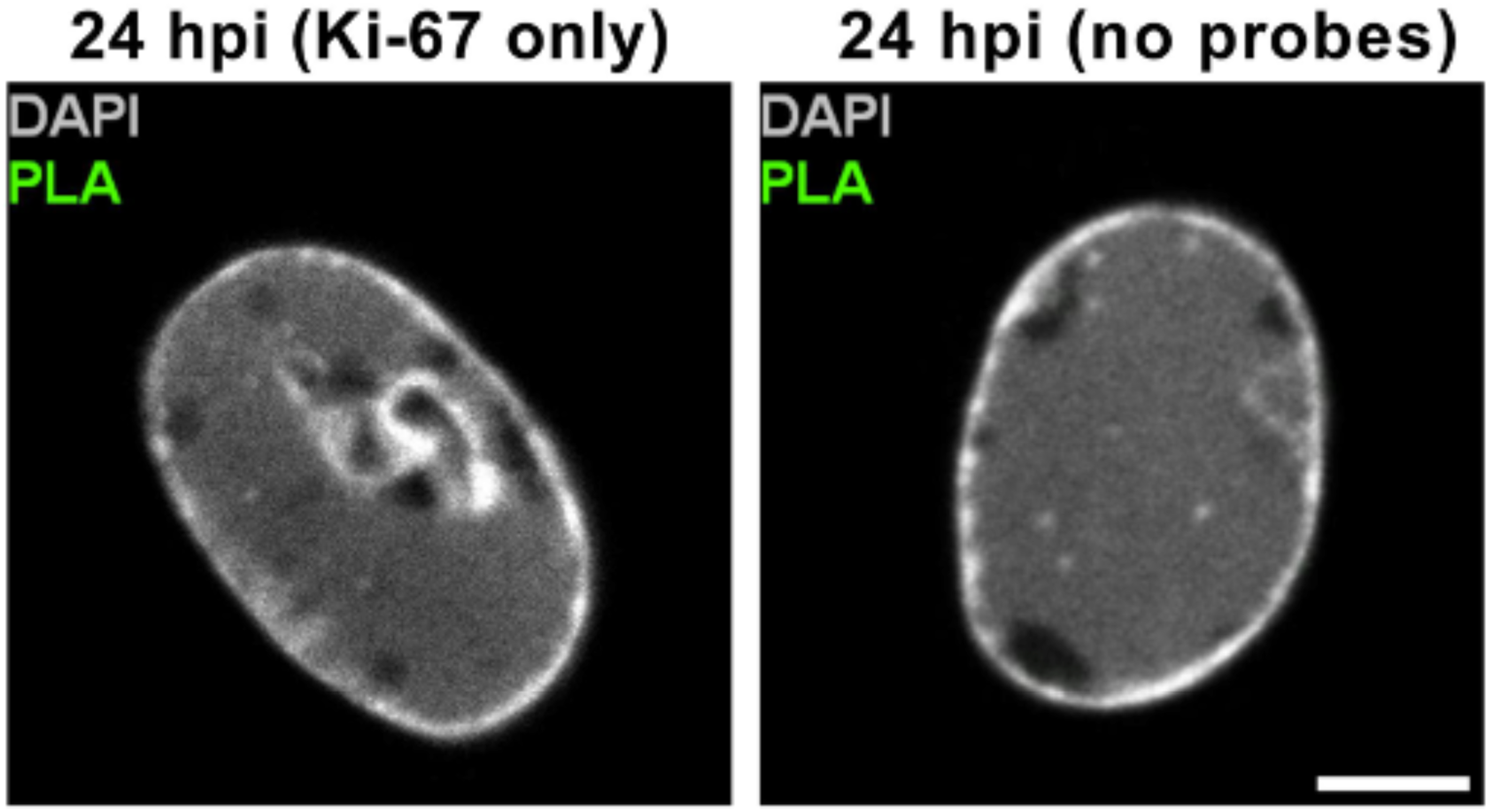
Technical controls for proximity ligation assay. Representative microscopy images of the technical controls for proximity ligation assay (PLA) are shown. Controls were performed with only Ki-67 antibody or without probes in 24 hpi infected cells. PLA foci in green and DAPI in grey. Scale bars, 5 µm.

**Figure S7.**
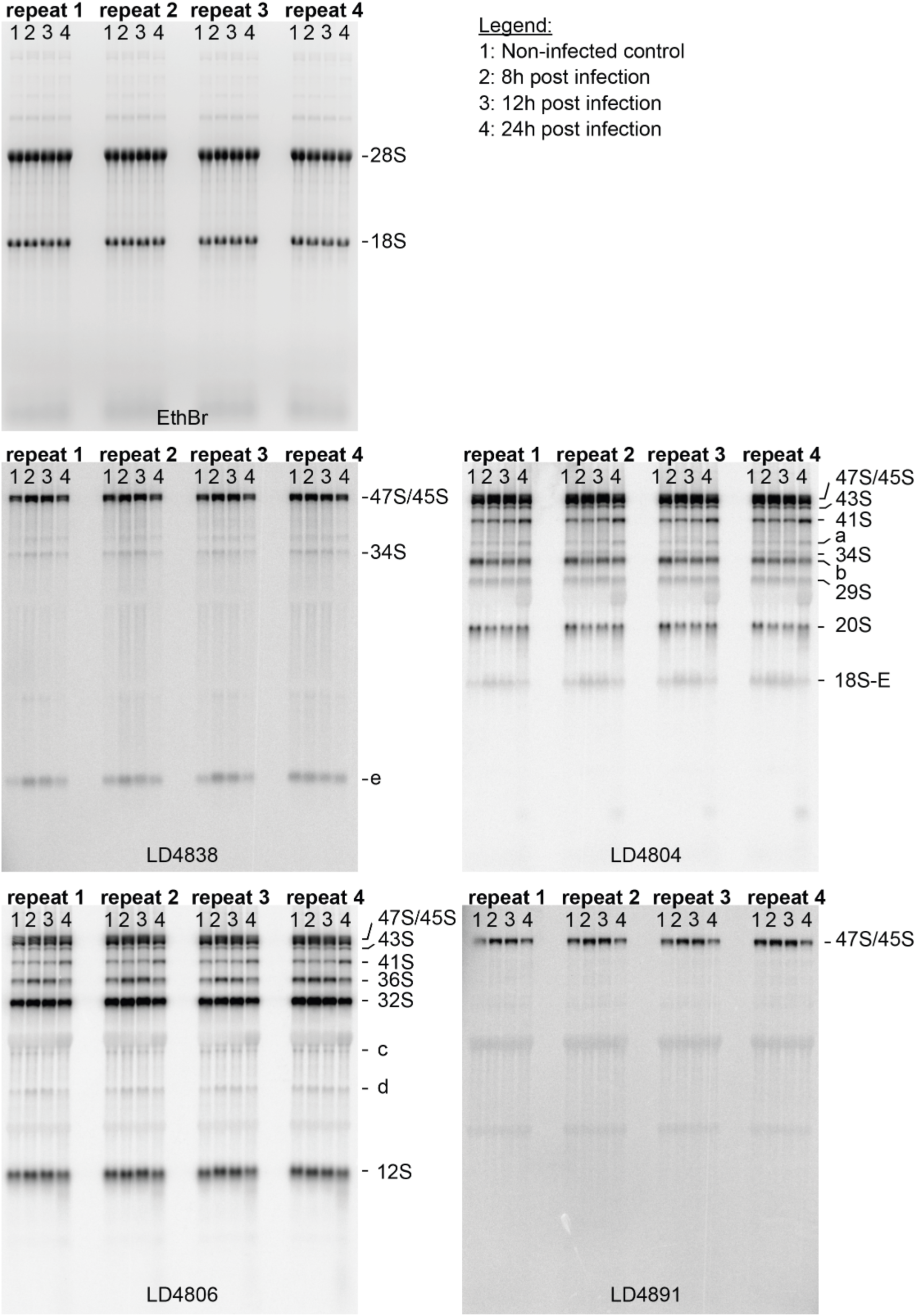
Pre-rRNA analysis of infected cells (4 independent biological samples) Agarose gel analysis of ethidium bromide-stained gel (a) and northern blotting (panel b-e, see Figure 7 legend for details of probes used). Total RNA was extracted at 8, 12, and 24 hpi. A noninfected (NI) sample was used as a control. The analysis of four independent biological replicates attests to the robustness of our observations.

**Table T1.**
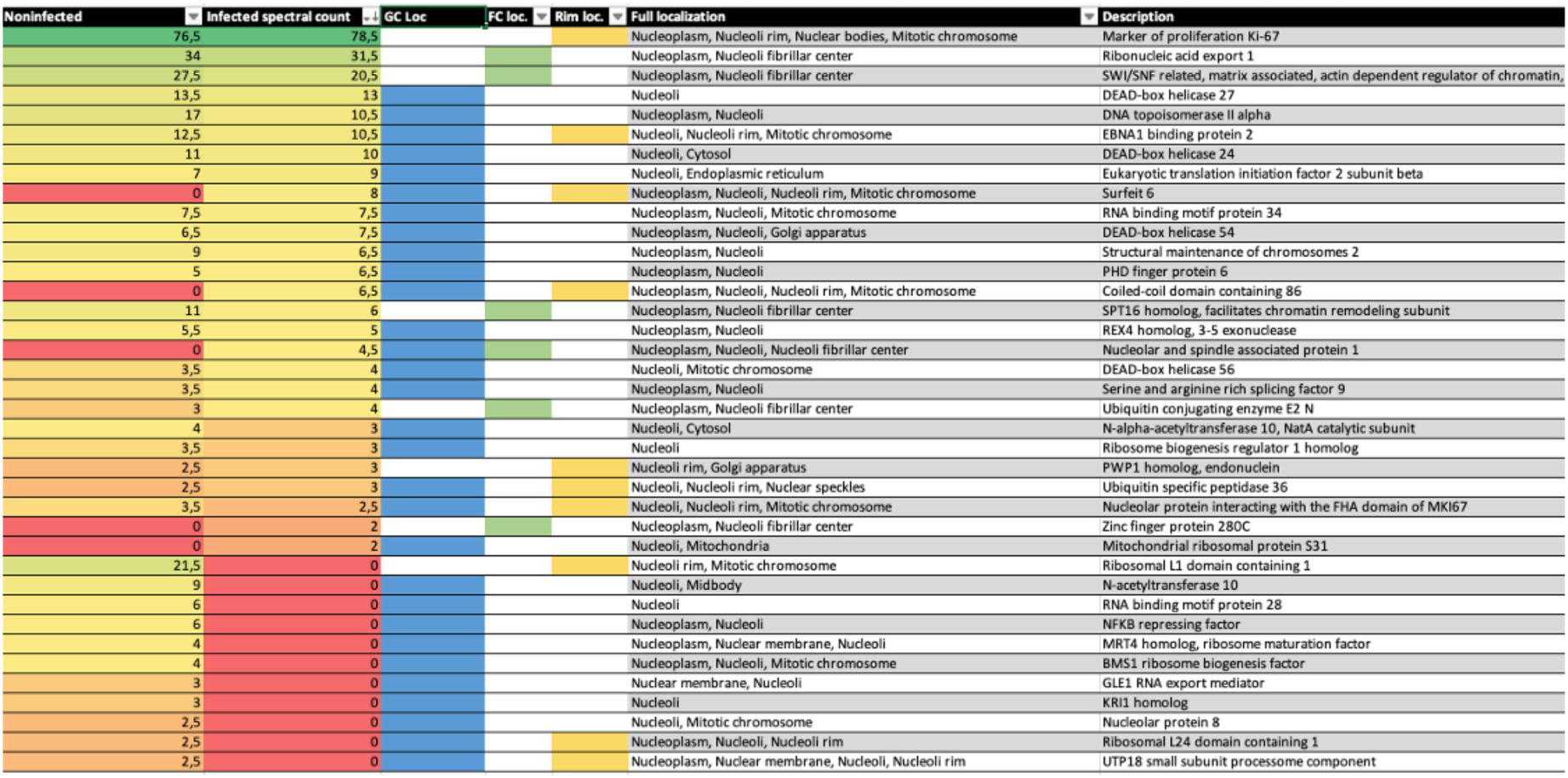
BioID hits linking NS2 to nucleolar organization and rRNA processing. The table presents 122 NS2-binding nucleolar proteins detected as high-confidence (BFDR ≤0.05) interactors. NS2-associated proteins localized in the fibrillar center (including FC and DFC, green), the granular component (GC, blue), and the nucleolar rim (NR, yellow) are highlighted.

**Table T2.**
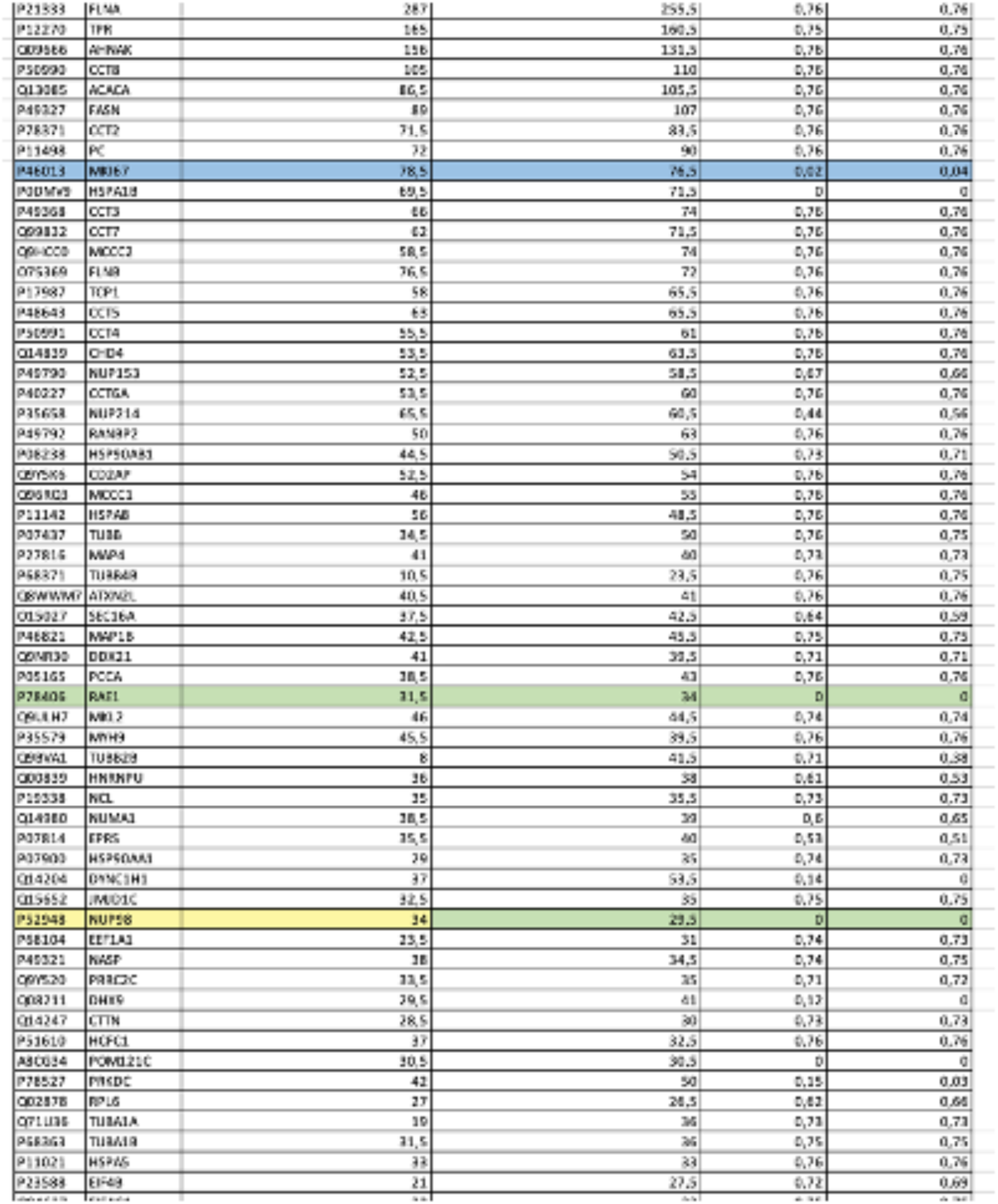
Unfiltered NS2 BioID data. Unfiltered SaintExpress-analyzed BioID data was used in this study. For each identified prey protein, the average spectral count in infected and noninfected samples and BFDR values used later for filtering the dataset are shown for each identified prey protein. NS2-associated proteins localized in the fibrillar center (including FC and DFC, green), the granular component (GC, blue), the nucleolar rim (NR, yellow), and dual localization are highlighted.

**Table T3.**
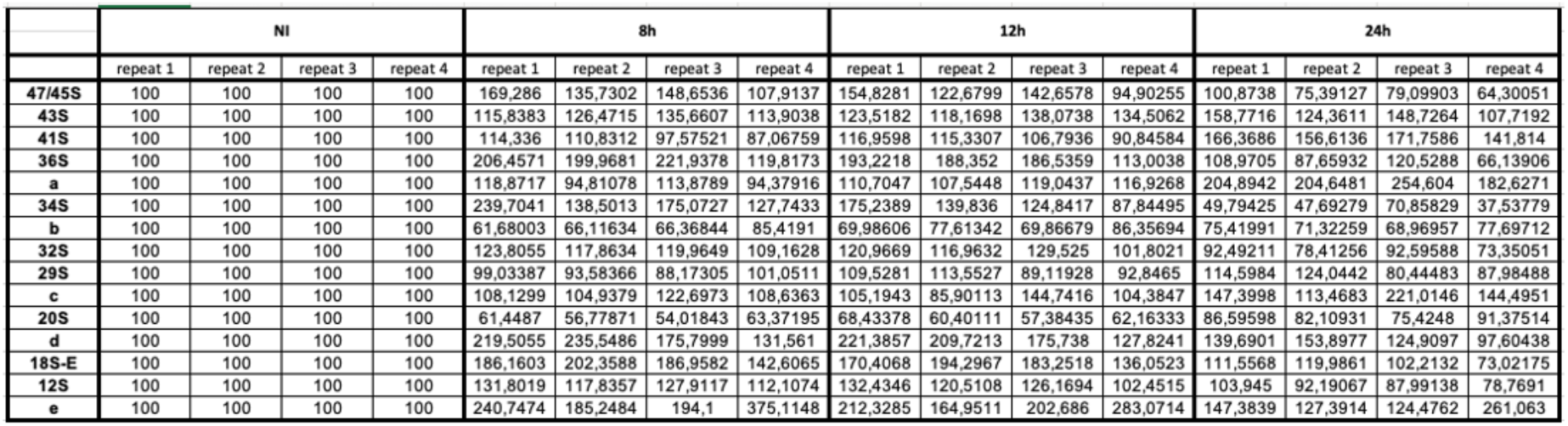
Quantification of rRNA processing in infection. Agarose gel analysis data of EthBr-stained RNA of noninfected (NI) and infected cells at 8, 12, and 24 hpi. The analysis of four independent biological replicates.

**Movie M1. Infection-induced alteration of nucleolar structure**

Expansion microscopy images and their 3D reconstructions of noninfected and infected NLFK cells at 24 hpi (see Figure 3). The surface rendering of the segmented whole nucleolus (grey), the low-intensity protein regions (green) based on the total protein staining, and DNA (blue) are shown. Specifically, interconnected low-intensity protein channel networks and infection-induced large spherical vacuolar cavities are visible.

**Movie M2. Structural changes in nucleoli of infected cells**

3D cryo SXT images and their reconstructions of noninfected and infected NLFK cells at 24 hpi (see Figure 4). The localization of the nucleus (blue) and nucleoli (green) are shown. Specifically, the number of small, low-density foci increases during infection.

